# Primary envelopment of Kaposi’s sarcoma-associated herpesvirus at the nucleoplasmic reticulum

**DOI:** 10.1101/2025.04.01.646635

**Authors:** Alexa Wilson, Neale D. Ridgway, Craig McCormick

## Abstract

Herpesvirus egress begins with primary envelopment of newly assembled capsids at the inner nuclear membrane (INM) facilitated by a viral nuclear egress complex (NEC). Primary envelopment has been observed at the peripheral INM as well as at structures known as nuclear infoldings. Nuclear infoldings resulting from invaginations of the INM are known as Type-I nucleoplasmic reticulum (NR), whereas infoldings of both INM and outer nuclear membrane (ONM) are known as Type-II NR. Here, we report that Kaposi’s sarcoma-associated herpesvirus (KSHV) reactivation from latency and lytic cycle progression correlates with increases in both types of NR invaginations, but primary envelopment was restricted to Type-I NR and the peripheral INM. Over a time-course of infection, DNA-containing KSHV C-capsids were frequently observed budding into nuclear infoldings contiguous with the INM and sparsely decorated with nuclear lamina, characteristic of Type-I NR. These Type-I NR structures co-localized with puncta containing CTP:phosphocholine cytidylyltransferase (CCTα), the enzyme that catalyzes the rate-limiting step in phosphatidylcholine (PtdCho) synthesis, and drives *de novo* membrane biogenesis and membrane curvature required for NR expansion; this may provide sufficient Type-I NR to match requirements for KSHV nuclear egress. The selective utilization of the Type-I NR for primary envelopment despite the concurrent expansion of the Type-II NR suggests a mechanism to target C-capsids into the Type-I NR. We also observed accumulation of de-enveloped KSHV C-capsids in 2^nd^ order nuclear infoldings, suggesting that primary envelopment and de-envelopment can occur not only at the nuclear periphery, but also at the Type-I NR.

**Importance:** Herpesvirus capsids assemble in the cell nucleus but are too large to exit via nuclear pores. Instead, they invade the inner nuclear membrane and acquire a provisional lipid envelope that is quickly shed through fusion with the outer nuclear membrane, a neat trick that delivers the capsid to the cytoplasm for subsequent steps in assembly and egress. Dynamic rearrangements of the nuclear membrane include invaginations that plumb the depths of the nucleus to facilitate cargo trafficking. Here, we demonstrate that during Kaposi’s sarcoma-associated herpesvirus (KSHV) replication, nuclear membrane infolding and expansion events increase, coinciding with recruitment of a host enzyme required for phosphatidyl choline synthesis to these sites. We observed an abundance of KSHV capsids at infoldings of the inner nuclear membrane, as well as free capsids in the interior of second order infoldings suggestive of de-envelopment events occurring prior to infold retraction. This complements a more well-established mechanism of KSHV egress at the nuclear periphery, and indeed, we observed concomitant egress of KSHV capsids via both mechanisms from the same nucleus. Similar observations for other herpesviruses by other groups suggests that this versatility in nuclear egress mechanisms is conserved.

## Introduction

Herpesvirus capsid assembly and DNA packaging occurs in a multistep process in the cell nucleus, yielding at least three distinct products known as A-, B- and C-capsids (1, 2). All capsid types possess a capsid shell, consisting of a major capsid protein (MCP) ORF25 (KSHV (3); homolog UL19 in herpes simplex virus 1 (HSV-1) (4)), small capsid protein ORF65 (KSHV (5); homolog UL35 in HSV-1 (6)), triplex proteins ORF26 and ORF62 which stabilize MCPs (KSHV (2); UL38 and UL18 in HSV-1 (7, 8)) and the portal protein ORF43 (KSHV; UL6 in HSV-1 (9)). A-capsids are considered abortive and constitute a capsid shell lacking internal scaffolding proteins and the viral genome. B-capsids represent an assembly intermediate or an abortive capsid and contain an inner scaffold consisting of scaffold protein ORF17.5 and scaffold protease ORF17 (KSHV(10); UL26 and UL26.5 in HSV-1, respectively(11)) required for genome loading, yet B-capsids do not contain the viral genome. C-capsids are mature capsids that have ejected scaffolding proteins and replaced them with the linear viral genome. There is evidence that C-capsids gain priority access to the host cell cytoplasm for subsequent steps in assembly and egress, although this quality control mechanism remains poorly characterized. Recently, a new capsid type termed ‘D-capsid’ was described; these capsids are similar to A-capsids in lack a genome, but in this case, they appear to have lost a packaged genome instead of failing to load one completely (12). Herpesvirus capsids assembled within the nucleus are ∼125 nm in size (2), and too large to fit through the nuclear pores through which the viral genome had entered, which can only accommodate up to ∼39 nm (13). Instead, herpesviruses bud through the inner nuclear membrane (INM) and into the peripheral nuclear membrane (NM) space, acquiring an envelope in the process. This event has been termed ‘primary envelopment’. Enveloped viruses in the perinuclear space fuse with the outer nuclear membrane (ONM), losing their envelope in the process of gaining access to the cytoplasm. Primary envelopment requires the nuclear egress complex (NEC) which consists of a type-II membrane anchored protein called ORF67 (KSHV (14); UL34 in HSV-1 (15), UL50 in human cytomegalovirus (HCMV) (16), BFRF1 in Epstein Barr virus (EBV)) and soluble protein ORF69 (KSHV; UL31 in HSV-1, UL53 in HCMV, BFLF2 in EBV). The NEC localizes to the INM where it induces local membrane curvature and recruits C-capsids for budding into the perinuclear space.

Although primary envelopment at the peripheral NM remains the central dogma of herpesvirus egress, mounting evidence suggests that this model does not accurately portray the process of primary envelopment. To date, the alpha-herpesviruses herpes simplex virus 1 (HSV-1) and pseudorabies virus (PRV), beta-herpesviruses human cytomegalovirus (HCMV), and the gamma-herpesvirus murine herpesvirus 68 (MHV-68) (17–20) have been shown to acquire their primary envelopes by budding into nuclear infoldings (NI). NI are similar in structure to the nucleoplasmic reticulum (NR) (reviewed in (5,10), Figure 1). The NR are classified as Type-I and Type-II based on membrane morphology and the contents of each nuclear invagination (Figure 1). Type-I NR are single membrane branched or unbranched invaginations of INM which extend into the nucleoplasm. Type-II NR are double membrane invaginations of both the inner and outer NM. As such, Type-II NR has nuclear pores and nuclear lamina, and facilitates an extended cytoplasmic interface with the nucleoplasm, while the Type-I NR is lamin-poor and devoid of proteins associated with the ONM.

**Figure 1.**
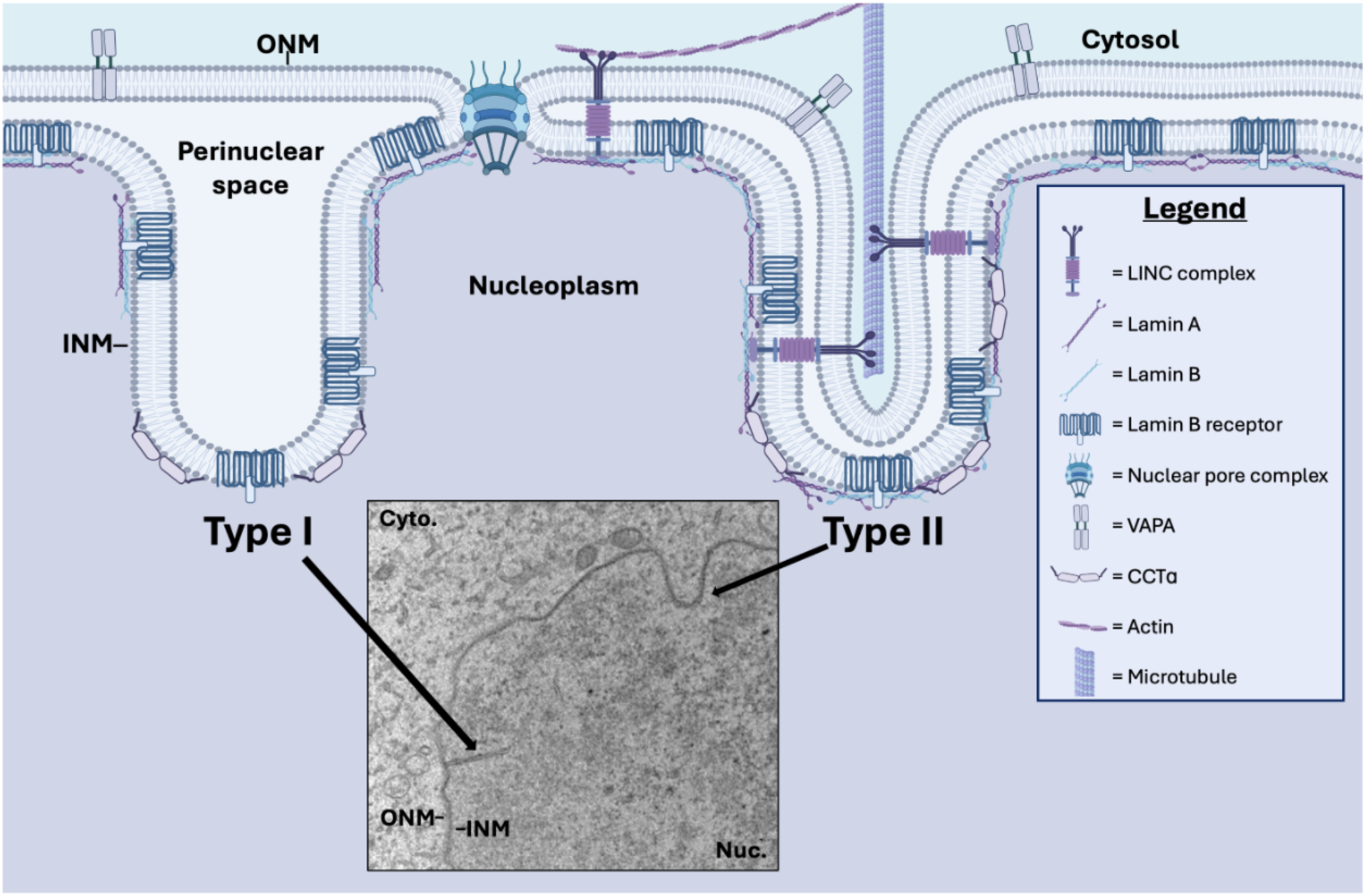
Morphology of Type-I and Type-II nucleoplasmic reticulum (NR). Model depicting the outer nuclear membrane (ONM) and inner nuclear membrane (INM), nucleoplasm (purple), and cytosol (blue). The inset shows a transmission electron microscopy (TEM) image of Type-I and Type-II NR in latent iSLK cells. Invagination of the INM yields the Type-I NR that is lamin poor and exclusively contains INM-associated proteins, such as the lamin B receptor. Invagination of the ONM and INM together produces Type-II NR with an underlying lamina and markers including: (i) lamin B receptor, (ii) vesicle-associated membrane protein-associated protein A (VAPA), which is involved in the delivery of late endosomes and the nuclear transfer of extracellular vesicle-derived components to the NR (49), (iii) the linker of nucleoskeleton and cytoskeleton (LINC) complex, which is tethered to the cytoskeleton and regulates NM dynamics, and (iv) nuclear pore complexes. Both Type-I NR and Type-II NR are decorated with CTP:phosphocholine cytidylyltransferase α (CCTα), an enzyme associated with the INM that catalyzes the rate-limiting step in synthesis of phosphatidylcholine for membrane biogenesis. CCTα and lamins cooperate to produce the NR (51).

It has been long recognized that CMV egress involves primary envelopment at NI. As early as 1964, intranuclear structures highly reminiscent of what we now refer to as NI were observed during murine cytomegalovirus (MCMV) infection (22). In the proceeding years, intranuclear compartments were documented with heightened detail during MCMV infection and their role in primary envelopment became evident (23). Eventually intranuclear structures were documented for HCMV (24) and later referred to as ‘pseudo-inclusions’ of the nucleus (25). Regardless of whether we refer to these structures as intranuclear inclusions, compartments, structures or pseudo-inclusions, a common theme from early ultrastructural analysis is that CMV buds into a distinct nuclear compartment during viral egress. More recent reports demonstrate that although NIs account for a mere 4.8% of nuclear membrane area during CMV infection, approximately 86% of nucleocapsids bud at NI rather than the peripheral nuclear envelope (18). NI appear lamin-poor, which could offer a route free of associated chromatin or nucleoli that would hinder CMV nucleocapsid budding (18). 3D analysis of NI during HCMV infection revealed a complex network of tubules and spherical compartments, with the latter assuming a variety of hierarchical structures (19). This network corresponds to the definition of Type-I NR from (26). It has been well-documented that lytic replication of herpesviruses causes the nucleus to increase in size (27). Villinger *et al.* proposed the ‘pushing membrane model’ to marry the changes in nuclear membrane size with the complex NR network observed during HCMV infection. In the pushing membrane model, HCMV causes a disruption of the nuclear lamina accompanied by the synthesis of new membrane material, resulting in the invagination of the INM into the nucleoplasm to form complex membrane structures. The resulting increase in nuclear membrane area enhances primary envelopment of nucleocapsids.

The alphaherpesvirus PRV buds into nuclear structures described as ‘tegusomes’, membrane structures apparently originating from cytoplasmic invaginations into the nucleus (17). Mature capsids bud into these structures, obtaining an envelope in the process. Tegusomes are sometimes connected with the NE and open into the cytoplasm, and most are comprised of single membrane with occasional double-membrane structures. Retrospectively reviewing TEM images of tegusomes reveals remarkable similarity to the Type-I and -II NR, as well as structures defined more recently by Buser *et al.,* and Villanger *et al.,* which we now know to be the Type-I NR. The PRV nuclear egress complex (NEC) consists of the inner nuclear membrane-anchored protein called pUL34 and soluble protein pUL31 (28). The PRV NEC induces local INM curvature and drives the formation of vesicle (29, 30) embedded within the Type-I NR (31, 32). Similarly, the NEC of the gammaherpesvirus MHV-68, consisting of ORF67 and ORF69, promotes nuclear invaginations positive for lamin A/C (33). Co-expression of ORF67 and ORF69 resulted in nuclear structures described as ‘vesicles wrapped by membranous structures in the nucleoplasm that appeared to have no connection to the nuclear membrane’ (33). During HSV-1 infection, the NEC proteins UL31 and UL34 localize to the INM and induce vesicle formation (34). The HSV-1 NEC promotes the reorganization of lamin, which acts as a structural barrier to capsids accessing the INM, causing lamin-positive protrusions in the intranuclear space that resemble the NR (35). Additional lamin modifications occur during herpesvirus infections. During HSV-1 infection the viral kinase US3 and a complex containing the NEC, the viral protein γ_1_34.5, host protein p32 and protein kinase C phosphorylate and destabilize the nuclear lamina (36–40). Additionally, kinase UL13 from human herpesvirus 2 (HSV-2) (41), kinase BGLF4 from Epstein Barr virus (EBV) (42), and recruitment of the HCMV kinase UL97 by the NEC (43, 44) all result in phosphorylation and disorganization of lamina. Since lamin disassembly is a key strategy used by herpesviruses during nuclear egress and is a prerequisite for NR formation, the disassembly of lamin during herpesvirus infection should be seen as a crucial step in herpesvirus remodeling of the NR.

Unlike the alpha- and beta-herpesviruses, the use of NIs by gamma-herpesviruses is largely uncharacterized. A double-membrane structure separated from the nuclear envelope was observed by transmission electron microscopy (TEM) (Naniima *et al.,* 2021) in cells infected with Kaposi’s sarcoma-associated herpesvirus (KSHV) but the authors do not comment on the nature of this invagination. In a murine model, MHV-68 infection produced invaginations of the INM that contained enveloped capsids (20). Less often, a secondary compartment is present within these invaginations that contains un-enveloped capsids. Ribosomes present in the secondary compartment suggest that the lumen of these compartments is derived from the cytoplasm, and the un-enveloped capsids within them have underwent de-envelopment. Secondary compartments were occasionally observed that harbor nuclear pore complexes and resemble the Type-II NR (26).

Here we report the first evidence of a human gamma-herpesvirus initiating primary envelopment at the Type-I NR. High-resolution TEM revealed KSHV capsids budding into the Type-I NR that was continuous with the INM with a lumen that resembled the NE periplasm. Our study used ∼200 high resolution TEM images to document these compartments during different stages of KSHV lytic replication. Unlike previous work, we capture NR expansion as early as 24 h post-reactivation, and describe a new order of NI hierachy, called the convoluted membrane NI that orginates from the Type-I NR. KSHV C-capsids bud out of the INM derived compartment and into a sub-compartment within the NR. Immunofluorescence (IF) microscopy positively identified these structures as the NR based on their association with known markers such as lamin A/C, CCTα and VAPA. Similar to the alpha- and betaherpesviruses, we propose that KSHV utilizes the Type-I NR for nuclear egress, and our data is highly reminiscent of the structures observed during HCMV egress (19). In conclusion, KSHV primary envelopment occurs at the Type-I NR by a mechanism that is conserved among multiple human herpesvirus subfamilies.

## Results

### Nuclear infolding increases during KSHV lytic replication

NI described in previous studies of herpesvirus infection have many of the characteristics of the Type-I NR (18, 19, 26). To study NI during KSHV infection we used inducible SLK (iSLK) cells harboring the doxycycline-inducible BAC16 KSHV genome, a well-established model system in which lytic replication is triggered by expression of the viral replication and transcription activator (RTA) protein (45). Transmission electron microscopy (TEM) of iSLK BAC16 cells reactivated for 72h revealed that NIs were continuous with the INM, initiating as narrow tubular ‘necks’ that expanded into large spheroid compartments containing enveloped C-capsids (Fig. 2A). We applied a previously established hierarchical system of NI categorization (19) to our studies of KSHV infection, whereby 1^st^ order NIs contain C-capsids within a single membrane compartment and 2^nd^ order NIs have non-enveloped C-capsid containing vesicles within a single membrane compartment. As previously reported (3), our findings suggest that the lumen of the 1^st^ order NI is continuous with the perinuclear space. The lumen of the 2^nd^ order NI is reminiscent of the nucleoplasm and hypothesized to originate from invaginations of nucleoplasm or cytoplasm into the first order NI. 3^rd^ order infoldings originate from separate compartments within 2^nd^ order infoldings and have a lumen that does not resemble the nucleoplasm, cytoplasm, or perinuclear space. We describe a novel ‘convoluted membrane’ NI, which appeared as a single membrane containing multiple encapsulated vesicles or membrane whorls (additional whorl images highlighted in Fig. S1).

**Figure 2.**
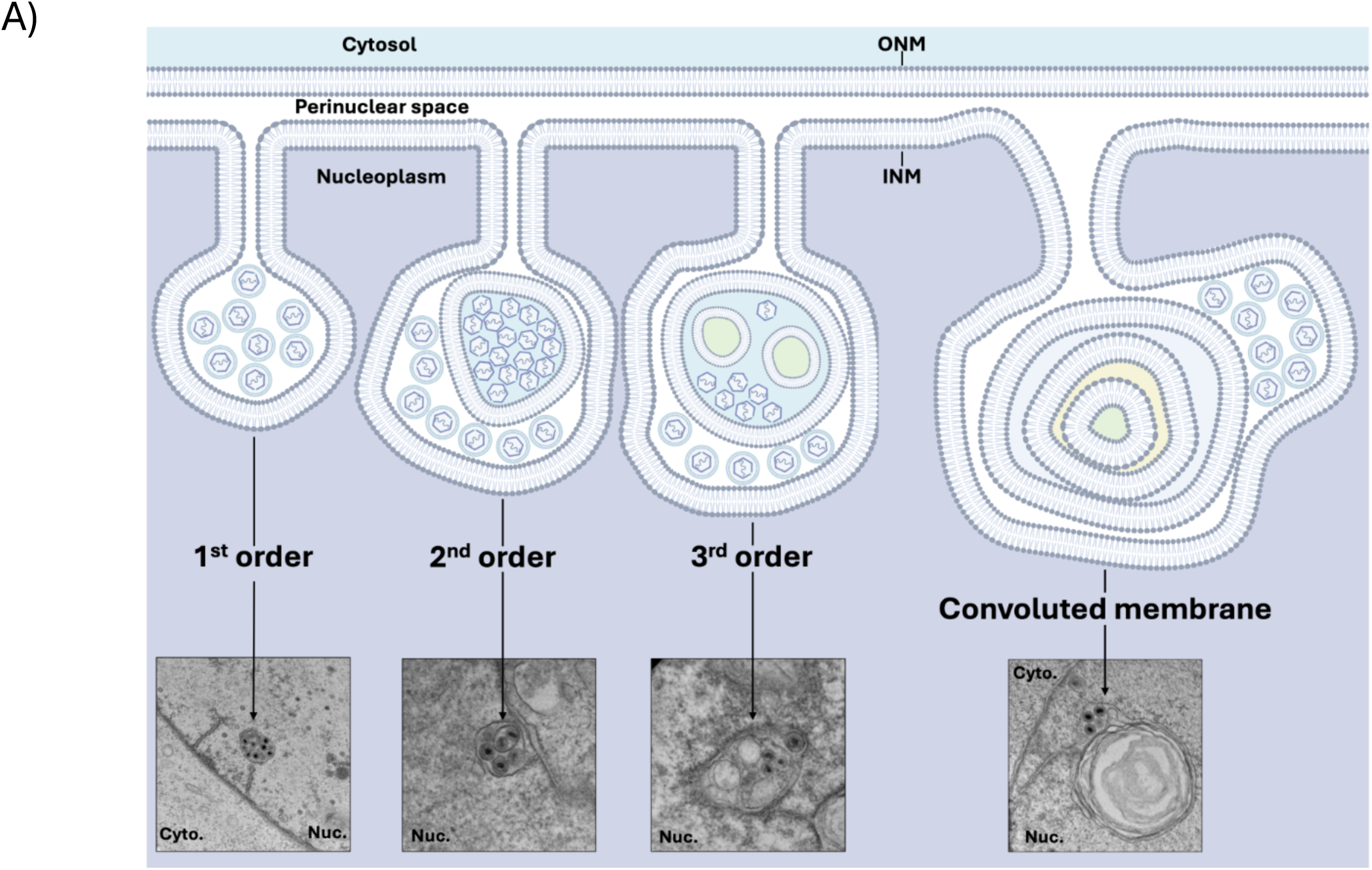
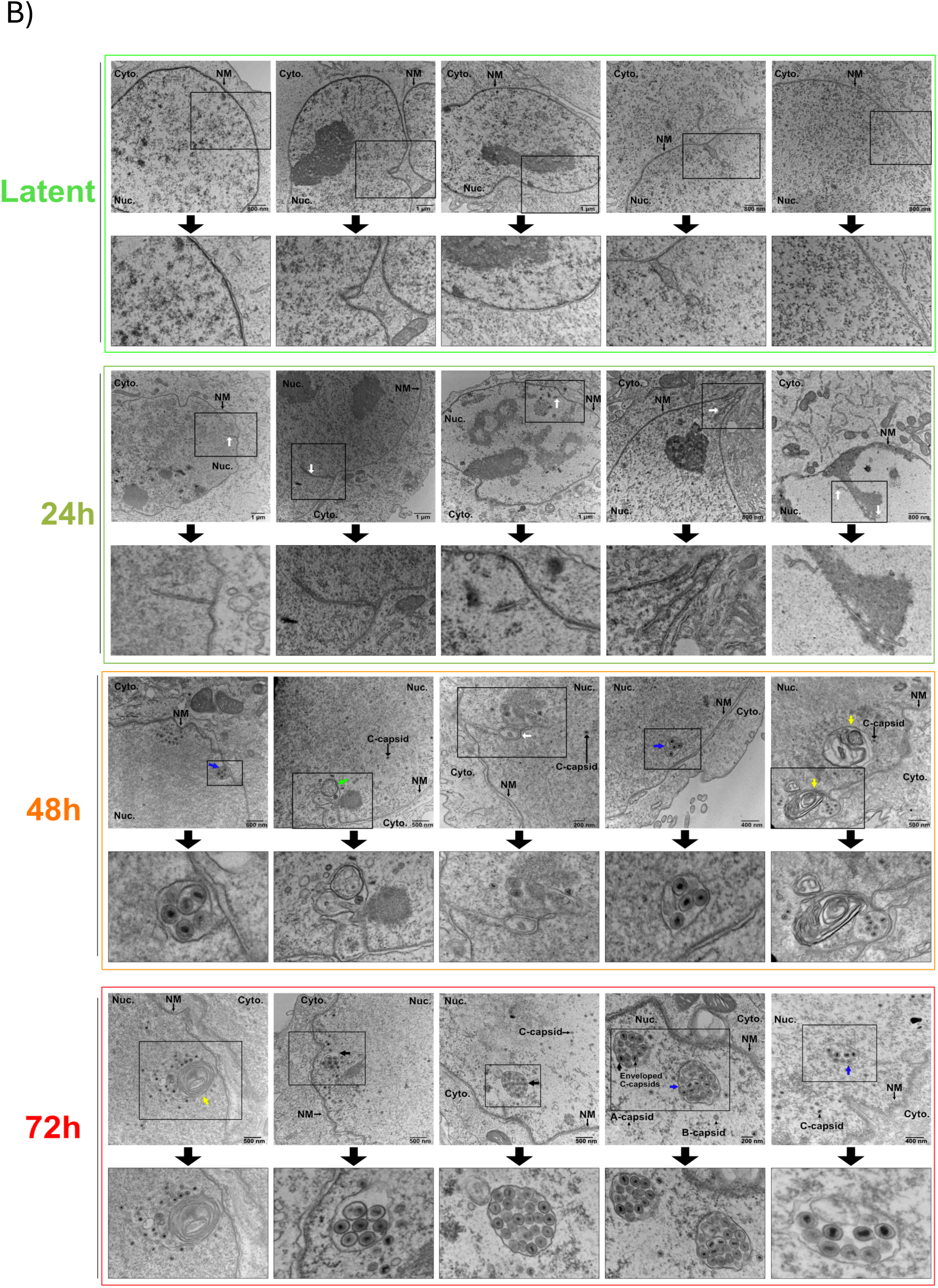
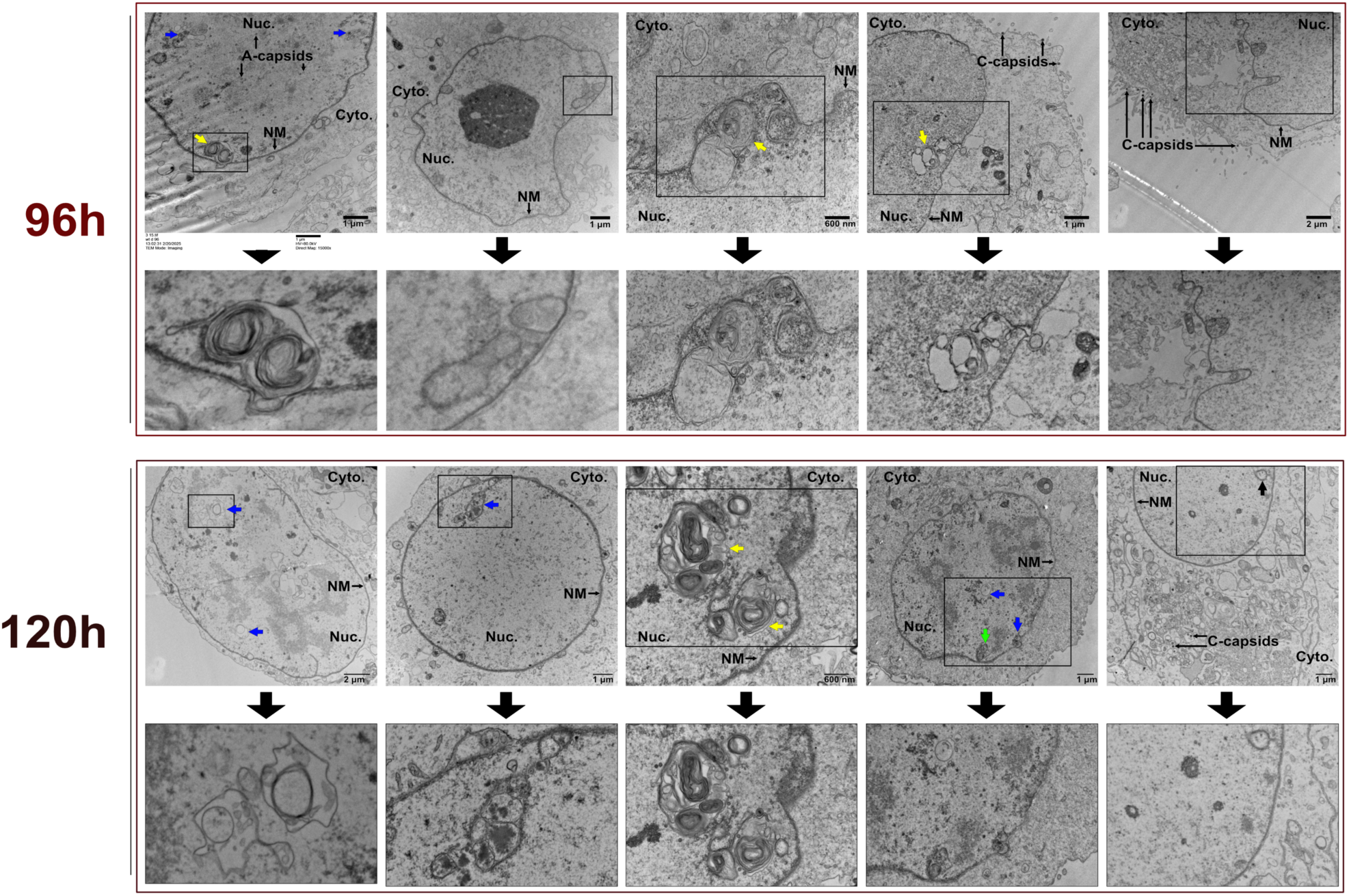
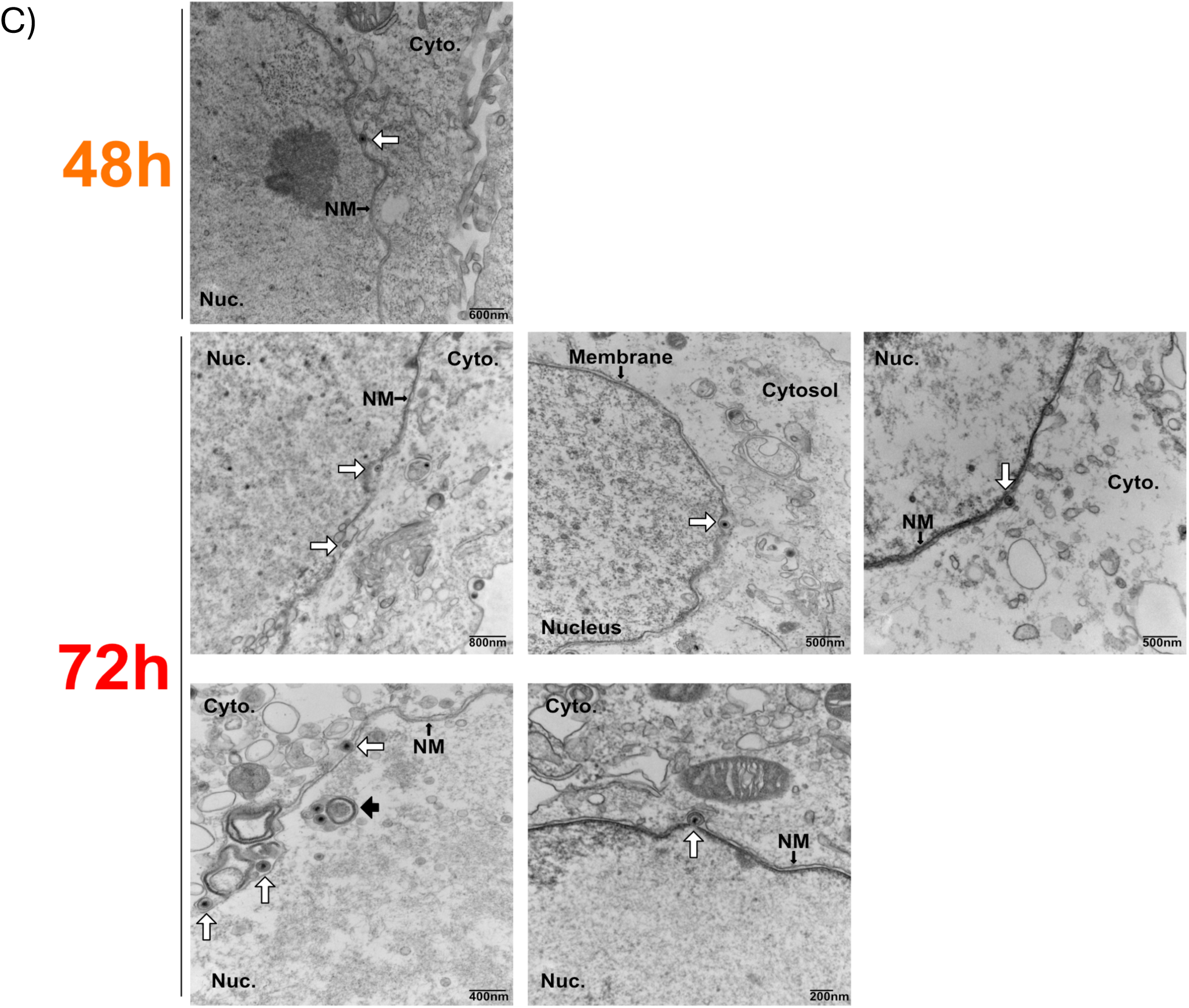
Enveloped KSHV C-capsids accumulate in diverse nuclear infoldings (NIs). (A) Orders of NIs observed during herpesvirus reactivation. The outer nuclear membrane (ONM) and inner nuclear membrane (INM) are shown in relation to the cytosol (blue) and nucleoplasm (purple). 1^st^ order NIs appear as single membrane tubules/vesicles that are continuous with the Type-I NR. 2^nd^ order NIs appear as a single membrane compartment that contains one or more additional single membrane compartments. 3^rd^ order NIs are 2^nd^ order NIs that contain additional compartments. Convoluted membranes often appear as a second order infold, but the second compartment is a large, convoluted membrane whorl. (B) Transmission electron microscopy (TEM) analysis of nuclear envelope remodeling in doxycycline-inducible iSLK cells infected with KSHV BAC16. Cells were reactivated with 1 μg/mL doxycycline and 1mM sodium butyrate and harvested at 24, 48-, 72-, 96 and 120-hours post-reactivation. Five representative images were selected per timepoint. Black boxes indicate areas of images enlarged below each parental image to allow NIs to be seen in detail. White arrows point to NIs. Black, blue, green and yellow arrows specifically point to 1^st^, 2^nd^, 3^rd^ and convoluted membrane infolds, respectively. Nucleus (Nuc.), cytoplasm (Cyto.) and nuclear membranes (NM) are labeled. (C) In addition to undergoing primary envelopment at NIs, C-capsids can still initiate classical budding at the peripheral nuclear envelope. TEM analysis of nine individual primary envelopment events at the peripheral INM are shown. In image 4, a 3^rd^ order nuclear infold is present (black arrow) near three classical budding events. All classical budding events are highlighted by white arrows.

To visualize the morphological changes to the nucleus over the course of viral replication, iSLK BAC16 cells were reactivated from latency and harvested at 24, 48, 72, 96, and 120h post-reactivation for TEM (Fig. 2B, additional images in Fig. S1). During latency, the nuclear envelope is smooth with evenly distributed lamin, and only a few small NIs are observed, indicating low basal NI activity similar to the basal activity seen in other human cell lines (46, 47). By 24h post-reactivation (early lytic replication), NIs increase in number, appearing as short tubules radiating from the nuclear periphery that are sometimes linked to spherical compartments. By 48h (late lytic replication), NIs expand into larger spherical vacuoles connected to the INM and often form networks. C-capsids enter 1^st^ order NI, acquiring a primary envelope, while some are unenveloped and present in 2^nd^ order NI. More complex 3rd order NI are also evident at this time. By 72h post-reactivation, NIs are more abundant and form large compartments containing numerous enveloped C-capsids. NIs appear as 1^st^, 2^nd^, 3^rd^ and convoluted membrane morphology. Strikingly, by 96h C-capsids have exited NIs, leaving behind empty pockmarked scars in cell nuclei. Scars appear as stretched out NI in which empty 1^st^ order infolds can be seen surrounding an empty 2^nd^ order infold. A few residual C-capsids speckle the nucleus at 96h; but most C-capsids are found in the cytoplasm or at the cell surface. Similarly, at 120h, empty pockmarked scars remain in cell nuclei, suggesting that these compartments are not repaired at this late stage of infection.

During KSHV lytic replication, NIs appeared near the nuclear periphery, as previously reported for HCMV lytic replication (18) (Fig. 2B). NIs containing enveloped KSHV C-capsids were often observed within single membrane compartments with 1^st^ order NI morphology (Fig. 2B, black arrows); however, C-capsids were also observed budding into a single membrane structure on the outside of a multi-membrane whorl (Fig. 2B, yellow arrows, Fig. S1). Additionally, a subset of NIs appeared to have a 2^nd^ order infold structure, and vesicles or tubules were seen inside a compartment akin to 1^st^ order NI (Fig. 2B, blue arrows). Although we captured significantly more capsids budding into NI, we still observed ‘classical’ budding at the peripheral NM at 72 h (Fig. 2C). As others have reported, these events are difficult to capture, perhaps due to the speed with which they occur (18). Given that we observed hundreds of instances in which C-capsids undergo primary envelopment at NIs (Supp. Fig. S1), compared to only eight budding events at the peripheral nuclear envelope (NE), it is likely that envelopment at NIs is preferentially utilized. We propose that NI-mediated budding represents an efficient mechanism that enables collective export of numerous capsids rather than one-by-one at the peripheral NE. This model is consistent with observations from HCMV infection studies, where hundreds of capsids have been reported to exit the nucleus *en masse* through similar structures (19). We observed a cell that contained three classical budding events at the peripheral NM, alongside a complex NI that contained multiple enveloped C-capsids within a 1^st^ order NI (Fig. 2C, black arrow). Thus, classical envelopment at the peripheral NM and envelopment at a NI can take place concurrently, both contributing to the transport of KSHV C-capsids to the cytoplasm.

### KSHV buds into Type-I nucleoplasmic reticulum

We observed accumulation of enveloped KSHV C-capsids in NIs that resemble the Type-I NR. However, definitive identification of these compartments as Type-I NR requires evidence of their direct connection to the INM. At 72 and 96 h post-reactivation from latency, we observed enveloped KSHV C-capsids within NIs that were clearly connected to the INM, but not the ONM, by a tubule (Fig. 3A, white arrows, blue colorization highlights continuity with the INM). Lamin extends up the tubule neck but is mostly absent around vesicles containing enveloped C-capsids (Fig. 3A, white dashed lines). Comparing the Type-I NR between 72 and 96 h, we observed that Type-I NR invaginations at 96h exhibit increased complexity and have a lumen with electron density consistent with nucleoplasm (Fig. 3A), similar to previous observations during lytic HCMV replication (19); similar observations were also made in MHV-68 lytic replication, but paradoxically, this lumen was thought to be of cytosolic origin (20). The lumen of the Type-I NR convoluted membrane is the same electron density as the lumen of the NE; we speculate that convoluted membranes result from hyper-proliferation of the INM during infection. An additional 13 images of Type-I NR that have a tubule protruding from them towards the INM are shown in Fig. S1. These observations are consistent with those of the Type-I NR, which contain a small amount of nuclear lamina where they attached to the INM but are elsewhere mostly devoid of lamina (21, 26). These observations agree with the Type-I NR observed during MCMV infection, which was also largely devoid of lamina (18). Of note, the Type-II NR also expands over the course of lytic replication (Supp. F2). Although we see C-capsids in close proximity to the Type-II NR at the cytoplasmic face of the nucleus, we have never observed a budding event at the INM of the Type-II NR. These results suggest that both the Type-I and Type-II NR expand during lytic replication, yet C-capsids specifically undergo primary envelopment at the Type-I NR.

**Figure 3.**
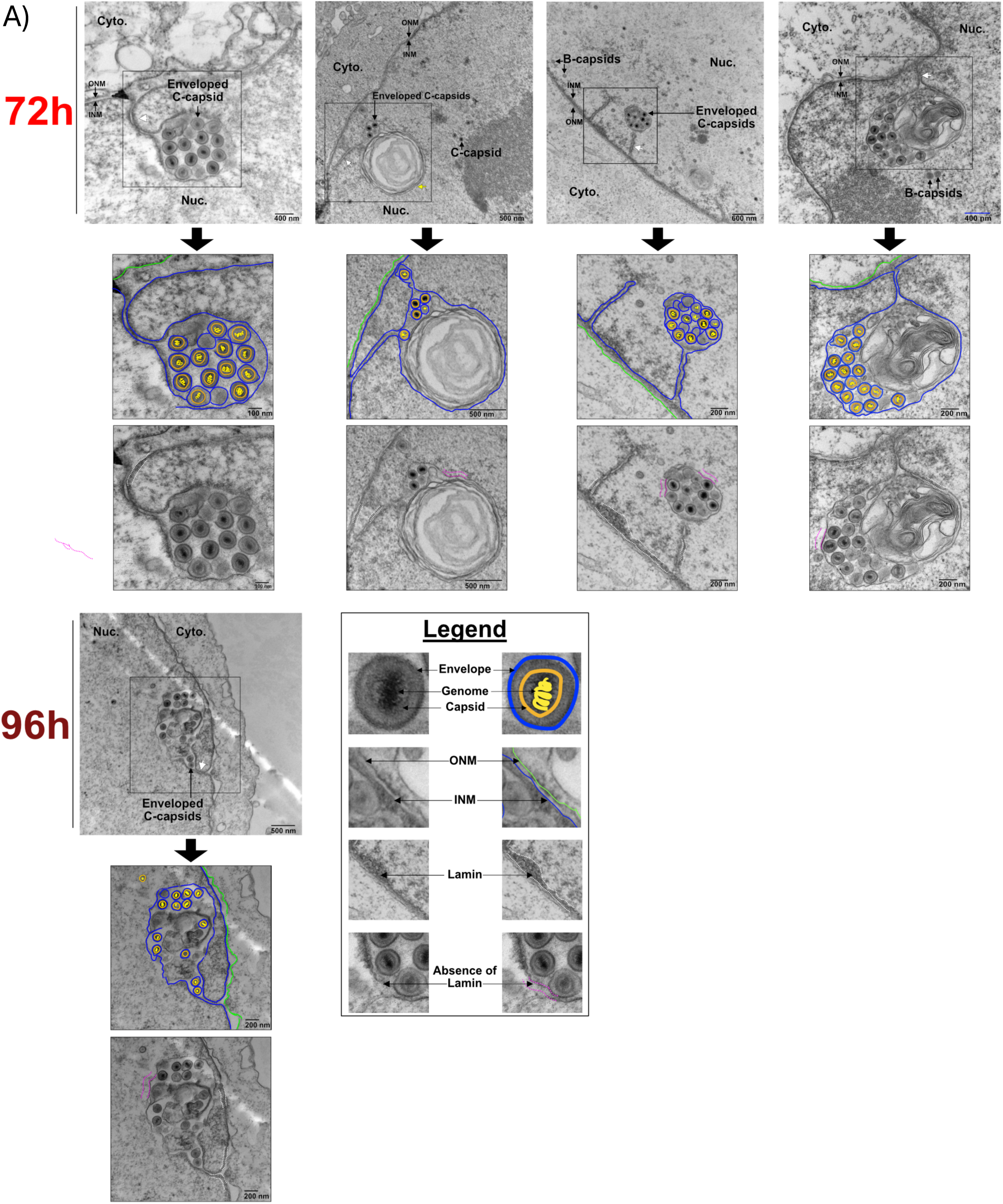
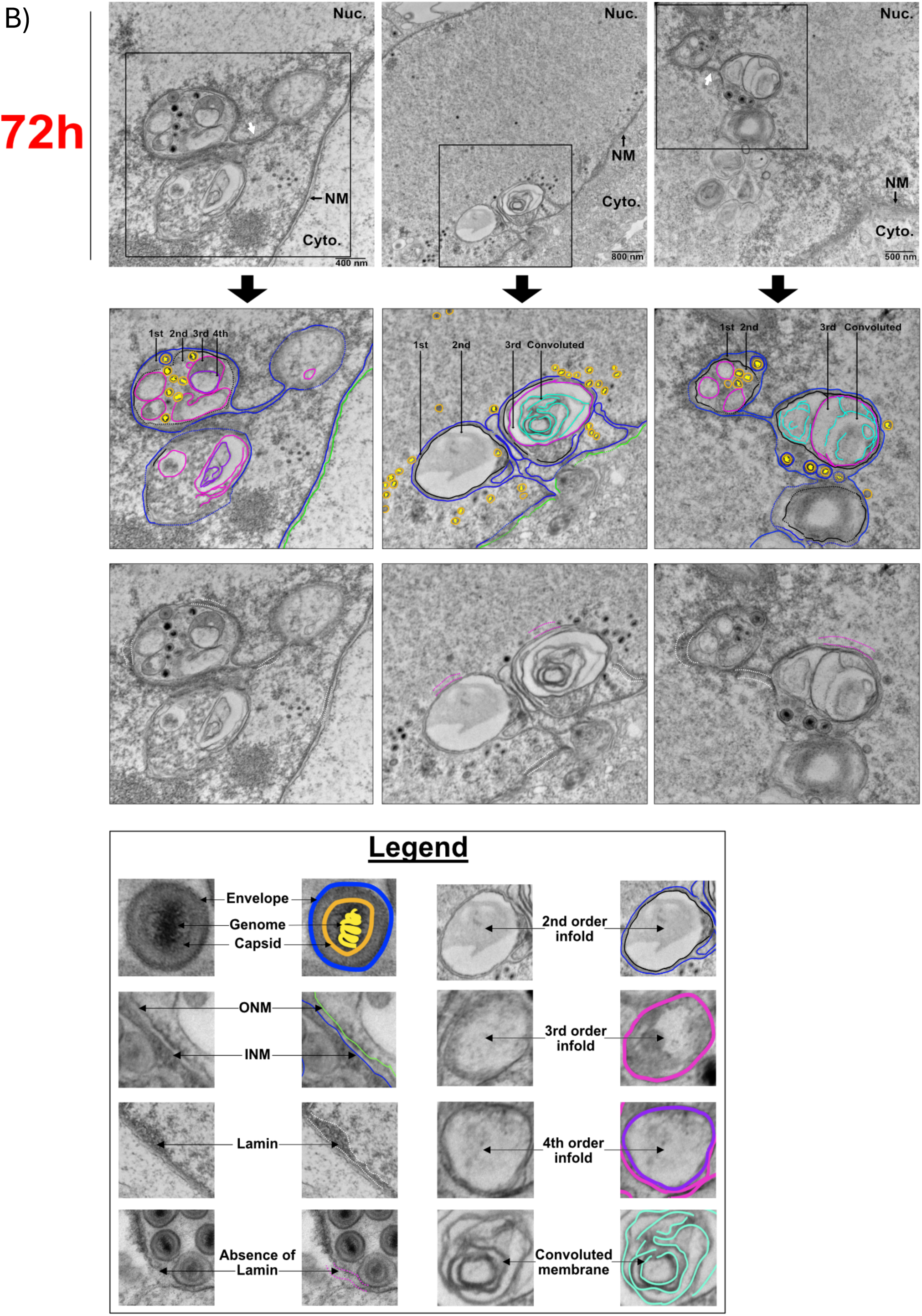
KSHV buds into Type-I nucleoplasmic reticulum. (A) Transmission electron microscopy (TEM) analysis of dox-inducible iSLK-BAC16 cells reactivated by the addition of 1 μg/mL doxycycline and 1mM sodium butyrate and harvested 72- or 96-hours post-reactivation. Lamin thickness is marked by white dashed lines for comparison across different NR regions. White arrows point to tubules connecting intranuclear compartments with the INM. Yellow arrow points to a convoluted membrane. (B) Type-I NR forms complex networks consisting of multiple spherical compartments connected by tubules. Compartment morphology varied, ranging between 1^st^, 2^nd^, 3^rd^ or convoluted membrane infolds connected by tubules. Enveloped C-capsids can be observed within 1^st^ order infolds, pertaining to the internuclear membrane space. Images in (A) and (B) are colorized to aid interpretation: INM (blue), ONM (green), enveloped C-capsids (blue), capsids (orange), and nucleoproteins (yellow).

During HCMV infection, Type-I NR is complex; consisting of a hierarchy of compartments interconnected by tubules (19). We documented numerous instances where spherical compartments containing enveloped C-capsids were connected to other spherical compartments by a tube shaped ‘stem’, resulting in the formation of a network (Fig. 3B; Fig. S1). Notably, we captured four C-capsids in the process of budding into the first order membrane of a Type-I spherical compartment in a single image (Fig. 3B; middle image). We frequently observed C-capsids budding into 1^st^ order NIs surrounding, and/or sometimes appearing to be contiguous with, convoluted membranes (Fig. S1), and yet only a single instance at 96h (Supp. F1) did one C-capsid access the convoluted membrane interior, indicating a potential barrier in the ability for capsids to access these membranes.

In conclusion, cross-sections through NIs during KSHV lytic replication demonstrate that they are continuous with the INM and match the definition of the Type-I NR (26). Similar to HCMV infection (19), the Type-I NR is remodeled into membranous networks during KSHV infection, characterized by multiple spherical compartments connected by tubule stems to the INM.

### Selective budding and fusion of C-capsids into Type-I nucleoplasmic reticulum

In iSLK-BAC16 cells harvested 72h post-reactivation TEM analysis revealed C-capsids in the process of budding into Type-I NR compartments, demonstrating that the enveloped C-capsids observed in earlier experiments access the Type-I NR by budding through the INM of these structures to obtain a primary envelope (Fig. 4A). A distinct C-capsid was observed at the periphery of the Type-I NR, partially enveloped by the INM, suggesting active budding into the NR (Fig. 4A; first image). The surrounding membrane appeared to conform tightly around this capsid, consistent with membrane curvature changes associated with capsid envelopment. This observation supports the role of the Type-I NR as a site of herpesvirus nuclear egress, facilitating capsid transit toward the cytoplasm. Notably, the NR displayed an expanded morphology, potentially accommodating multiple capsids undergoing this process concurrently or in quick succession. These findings align with previous studies in HCMV demonstrating that nuclear budding at the Type-I NR serves as a critical step for capsid transport through the nuclear envelope (18, 19).

**Figure 4.**
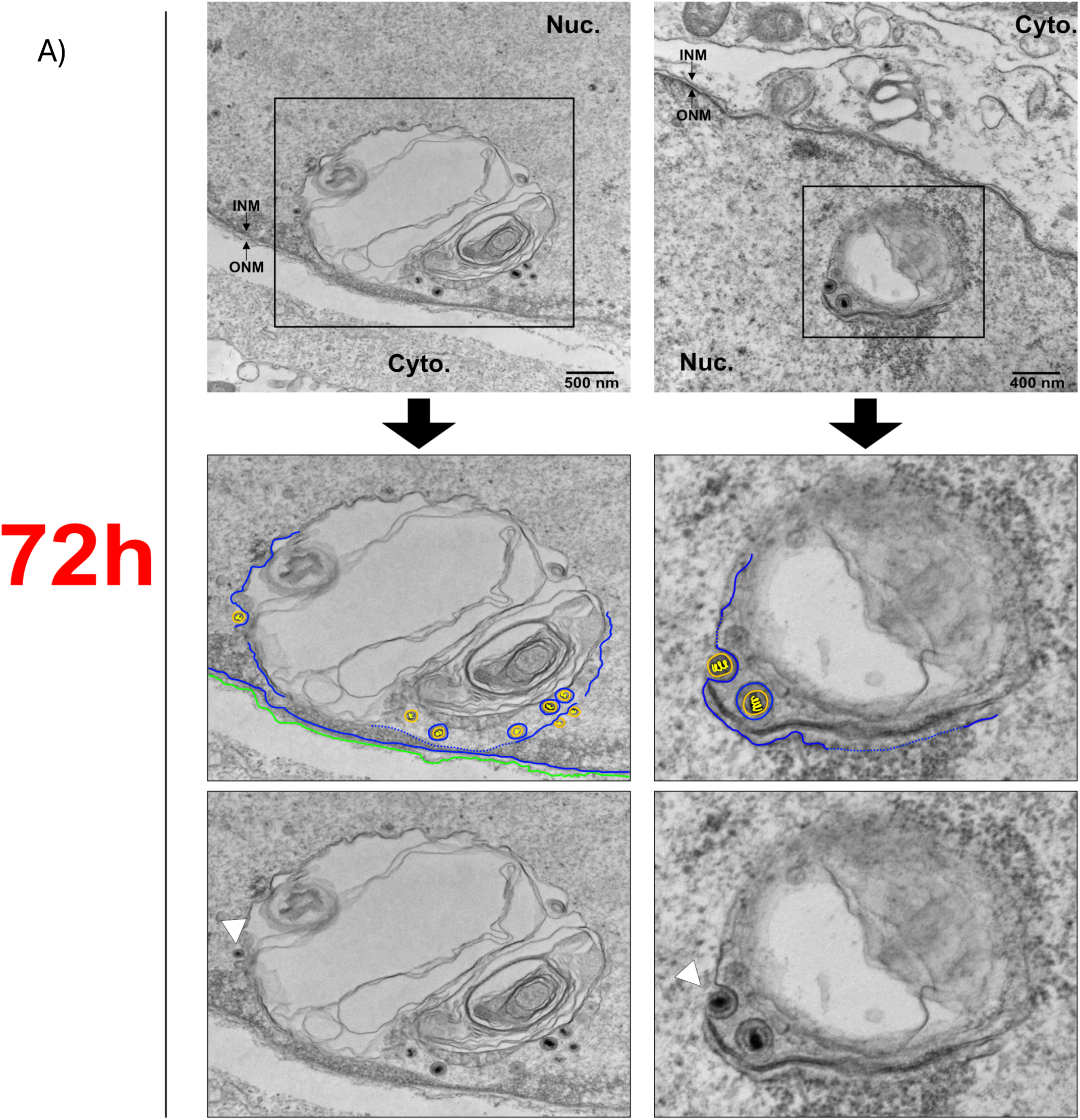
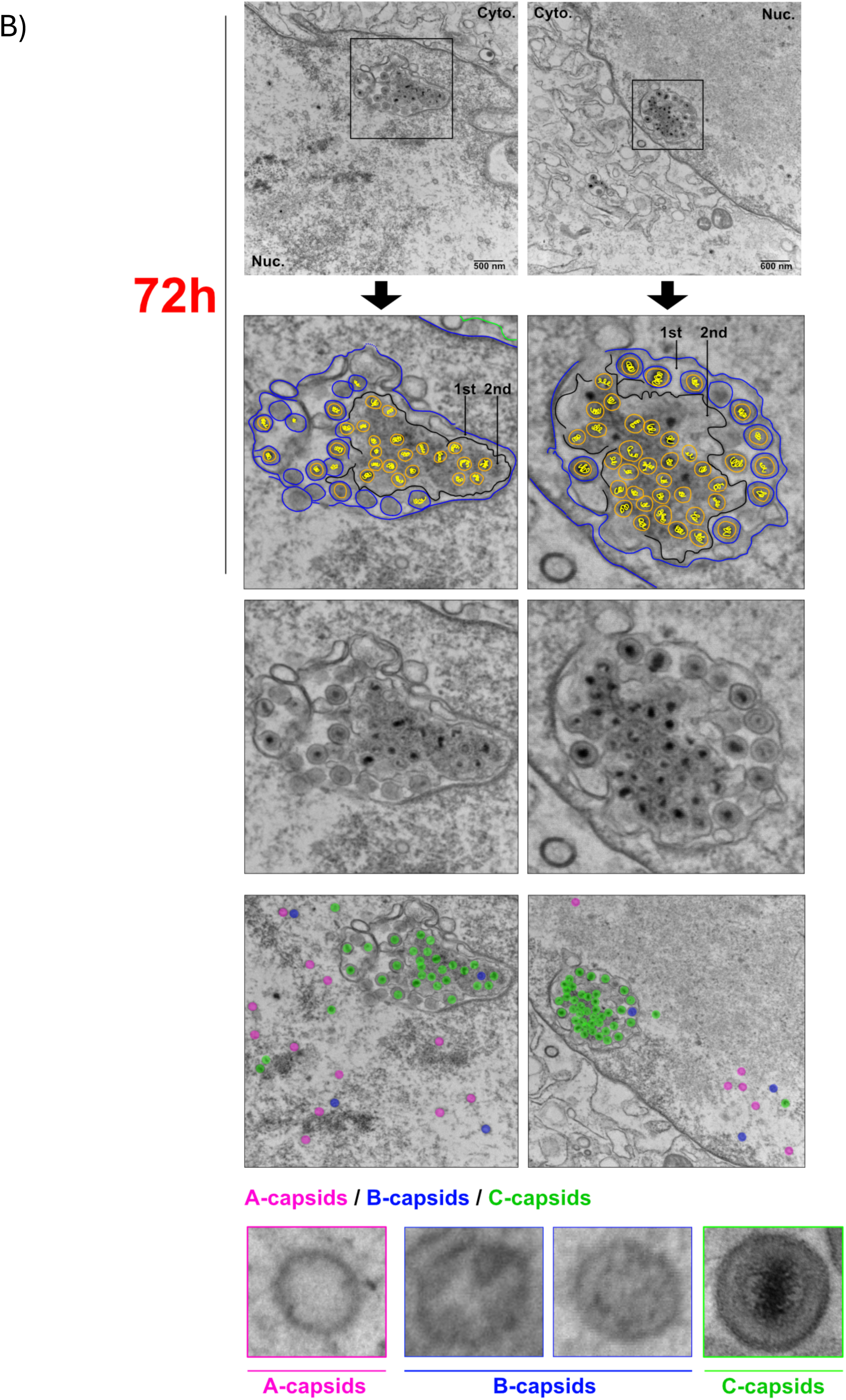
Selective budding and fusion of C-capsids into Type-I nucleoplasmic reticulum. (A) Dox-inducible iSLK-BAC16 cells were reactivated from latency and harvested 72h post-reactivation. Cell pellets were counterstained with osmium tetroxide and uranyl acetate, then processed for TEM. White tailless arrows show C-capsids budding into the Type-I NR. (B) Dox-inducible iSLK-BAC16 cells were harvested for TEM as described in (A). Enveloped C-capsids exit the NR, losing their envelope in the process. Unenveloped C-capsids accumulate within the 2^nd^ order NI.

Herpesviruses produce A-, B- and C-capsids (1). A-capsids do not contain the inner protein scaffold that supports viral genome loading and appear as an empty capsid shell (Fig. 4B). B-capsids contain the inner capsid scaffold and may display a morphology termed the ‘soccer ball’ capsid whereby the internal scaffold structure composed of ORF17.5 has been cleaved by the scaffold protease ORF17 and collapsed into itself (proposed to be either an intermediate of C-capsid assembly, or failed byproducts of capsid maturation(48)), or a concentric circle morphology where the major capsid protein has been processed (Fig. 4B). C-capsids are the mature, infectious capsids that contain the packaged viral genome within the capsid shell (dark black core in a circle, Fig. 4B). The Type-I NR was densely populated with C-capsids (Fig. 4B), indicating that these regions serve as major sites of viral egress, similar to observations during HCMV infection (19). The accumulation of C-capsids in these invaginations suggests a preferential localization of the NEC at these structures, likely facilitating efficient capsid envelopment and transport. The presence of multiple capsids in the NR further supports the hypothesis that the Type-I NR functions as specialized subdomain that supports coordinated bulk nuclear egress of KSHV capsids. These findings highlight the spatial organization of viral egress and suggest that the NEC may be concentrated at the Type-I NR to coordinate capsid envelopment and transport. Moreover, we observed a dynamic process in which enveloped C-capsids bud from the first order Type-I NR into the 2^nd^ order Type-I NR resulting in a loss of their primary envelope derived from the INM (Fig. 4B). The presence of multiple segregated enveloped and non-enveloped capsids within these infoldings indicates a regulated mechanism, potentially mediated by the NEC. This observed loss of the primary envelope is consistent with published models proposing that nuclear membrane fusion events occur within the NR, allowing C-capsids to be repositioned for subsequent transport out of the nucleus (19). Capsids from all TEM images included in this study were counted and their abundance in the 1) nucleoplasm, 2) nucleoplasmic reticulum, and 3) peripheral nuclear membrane space, were quantified (Table 1). At 48 hours post-reactivation, 18% of nuclear capsids were associated with the NR compartment, increasing to a peak of 30% by 72 hours. The majority of NR-associated capsids at all timepoints were localized to 1st order infolds, suggesting preferential capsid accumulation in the innermost NR structures. By 96 hours, the proportion of NR-associated capsids sharply declined to 3%, consistent with capsid departure from the 2^nd^ order infolds into the cytosol. Characteristic of KSHV infection, A-, B-, and C-capsids begin to accumulate around 48h sustained into 72h, when C-capsids are the most abundant capsid type in the nucleoplasm. By 96h, C-capsids have majority exited the nucleus leaving behind greater A- and B-capsid levels. These findings provide structural evidence for a multi-step nuclear egress pathway that specifically selects for C-capsids, where the hierarchy of NR architecture plays a critical role in herpesvirus capsid transit.

**Table 1.** Quantification of capsids within the nuclei of iSLK-BAC16 cells at 48-, 72-, 96-, and 120-hours post-reactivation. Total nuclear capsids were counted from all TEM images included in this study. Capsids were categorized as follows: (1) capsids associated with the nuclear replication (NR) compartment, further subdivided into (1.1) capsids located in 1st order infolds and (1.2) capsids located in 2nd order infolds, and (2) capsids associated with the peripheral nuclear envelope. At 48 hours post-reactivation, 18% of nuclear capsids were associated with the NR, reaching a peak of 30% at 72 hours. Most NR-associated capsids were consistently observed in 1st order infolds across all timepoints. By 96 hours, NR-associated capsids declined sharply to 3%, suggesting a temporal shift in capsid localization consistent with nuclear egress. While absolute capsid counts cannot be directly compared across timepoints due to differences in image sampling (with more images acquired at 48 and 72 hours), the proportion of NR-associated capsids within each timepoint reflects consistent and biologically relevant trends. Similarly, the distribution of A-, B-, and C-capsids at each timepoint aligns with established models of KSHV capsid maturation, further supporting the biological relevance of the observed temporal dynamics.

|  | 48 h |  |  | 72 h |  |  | 96 h |  |  | 120 h |  |  |
| --- | --- | --- | --- | --- | --- | --- | --- | --- | --- | --- | --- | --- |
| **Total capsids in NR** | 65 |  |  | 442 |  |  | 37 |  |  | 1 |  |  |
| Enveloped 1<sup>st</sup> order | 53 |  |  | 309 |  |  | 27 |  |  | 1 |  |  |
| Non-enveloped 2<sup>nd</sup> order | 12 |  |  | 133 |  |  | 10 |  |  | 0 |  |  |
| Peripheral NM capsids | 1 |  |  | 8 |  |  | 0 |  |  | 0 |  |  |
| **Capsid type** | A | B | C | A | B | C | A | B | C | A | B | C |
| Nucleoplasm | 75 | 63 | 214 | 139 | 414 | 939 | 203 | 861 | 270 | 11 | 136 | 50 |
|  | 21% | 18% | 61% | 9% | 28% | 63% | 15% | 65% | 20% | 6% | 69% | 25% |
| Total capsids in nucleoplasm | 352 |  |  | 1492 |  |  | 1334 |  |  | 197 |  |  |

### Type-I nucleoplasmic reticulum expansion in KSHV infected cells correlates with recruitment of CTP:phosphocholine cytidylyltransferase

We next conducted immunofluorescence microscopy studies on KSHV infected cells to determine the subcellular localization of key proteins involved in NR expansion, lamin A/C, VAPA, and CCTα (49). <u>V</u>APA engages <u>o</u>xysterol-binding protein–related protein 3 and <u>R</u>ab7 to form the <u>VOR</u> complex responsible for the transfer of extracellular vesicle (EV)-derived components from endosomes into the NR. This mechanism delivers nucleic acids and potentially other signaling molecules from internalized EVs to specific regions in the nucleoplasm, often near the nucleolus (49). CCTα is activated by insertion of its lipid sensing amphipathic helix into the INM where it catalyzes the rate-limiting step for phosphatidylcholine (PtdCho) synthesis required for *de novo* membrane biogenesis (50). CCTα also aids in NR expansion by promoting membrane curvature in collaboration with a lamin A/B1 scaffold (Fig. 1). Since the ER is continuous with the ONM, VAPA marks Type-II NR, whereas CCTα is associated with Type-I NR.

In latently KSHV infected cells, lamin A/C is uniformly distributed across the INM and intensified at the nuclear envelope edges (Fig. 5A). By 72h post-reactivation from latency, lamin A/C projections radiate from the INM, forming spherical compartments, consistent with NR expansion reported in previous studies of the NR (51, 52) and lamin reorganization during HCMV, EBV, HSV-1 and HSV-2 infection (42, 53–55). By 96h, lamin A/C-positive tubules emerged from the INM, exhibiting a branched, cylindrical morphology and linking multiple NR compartments. One limitation of this lamin A/C immunostaining procedure is that it could not detect lamin-poor NR compartments that were readily detected by TEM (Figs. 2B, 3A-B, 4A-B); instead, these immunostaining images showed lamin A/C positive tubules lacking a terminal compartment, which is likely undetectable because it is lamin-poor.

**Figure 5.**
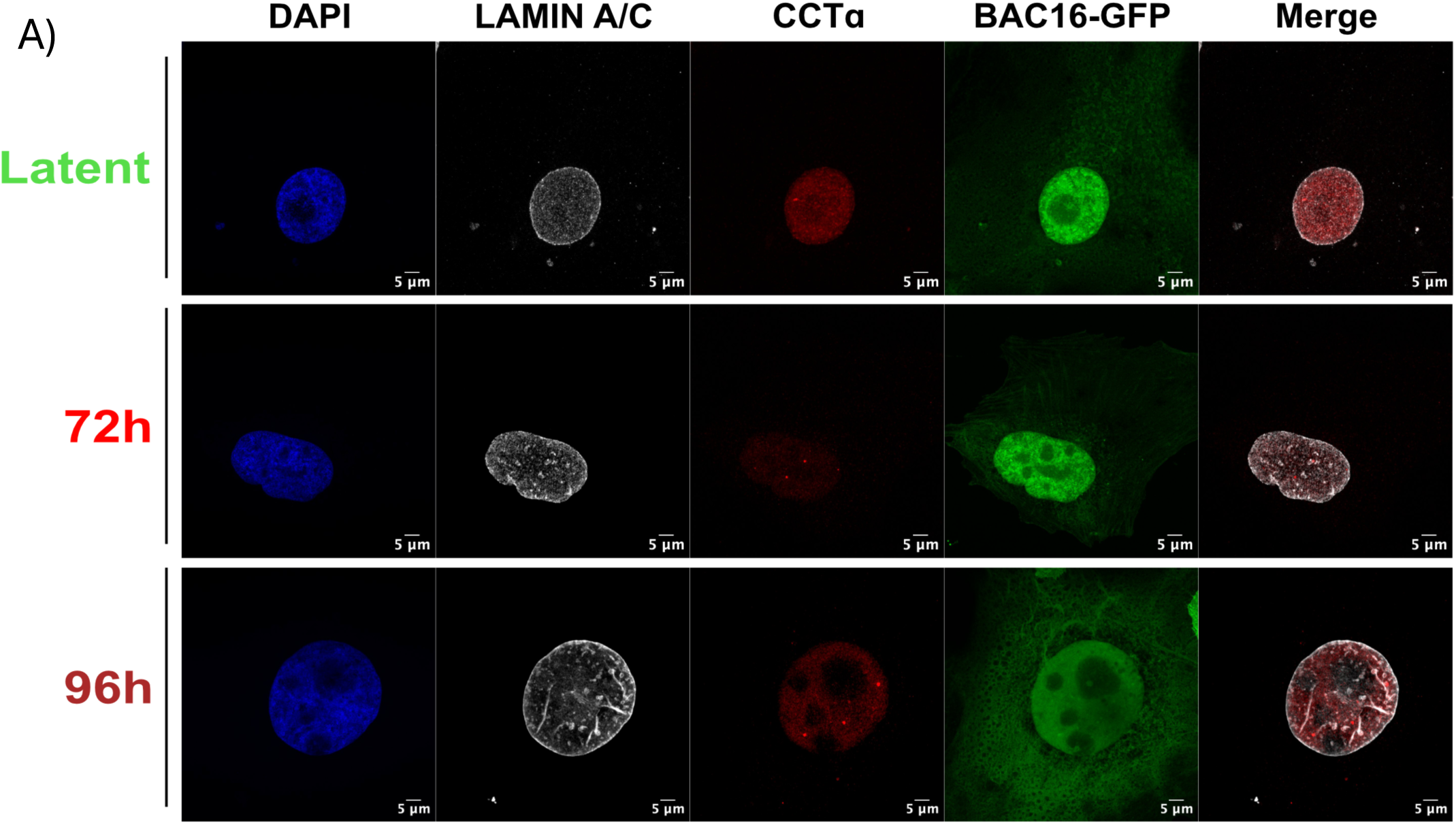
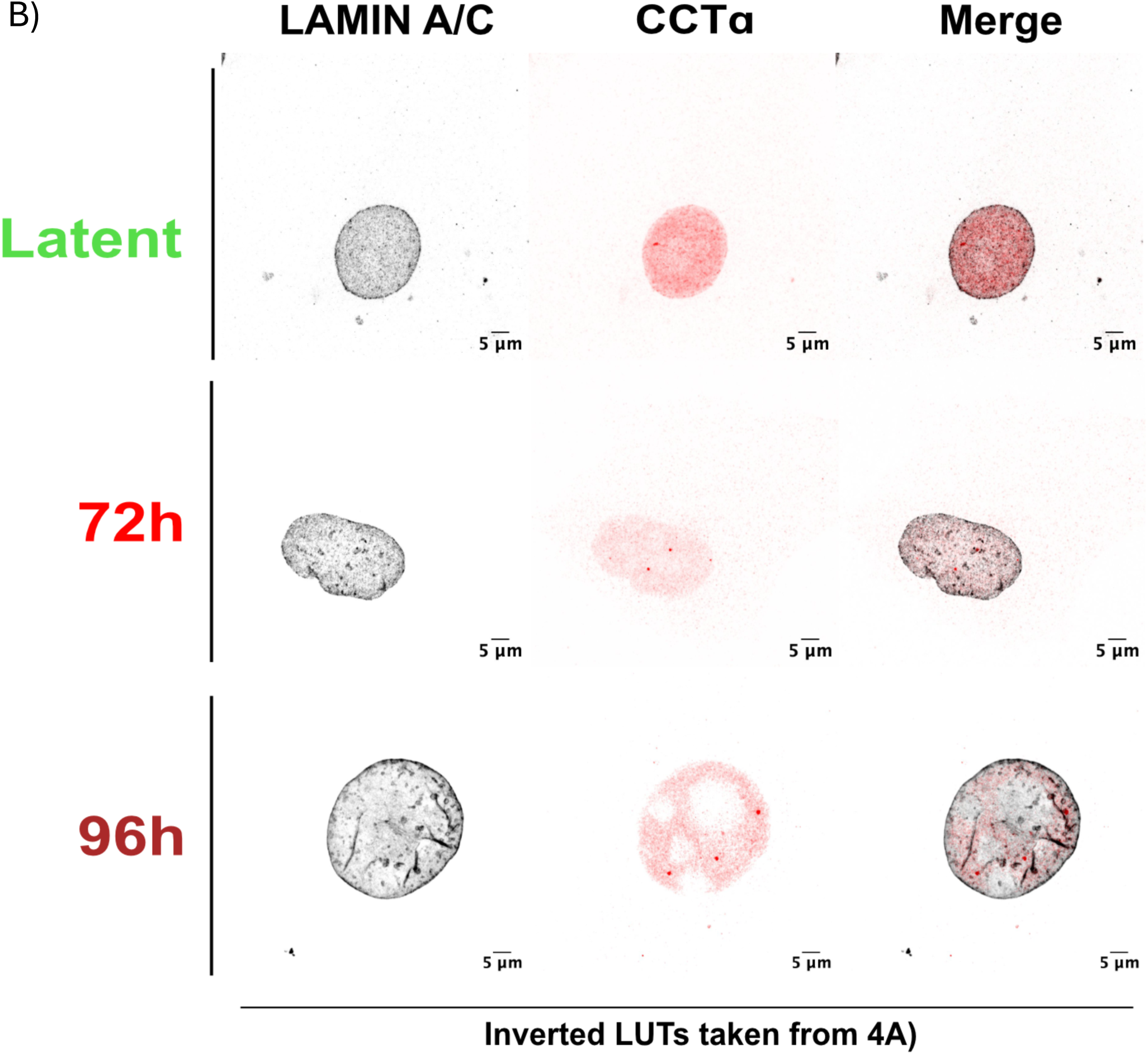
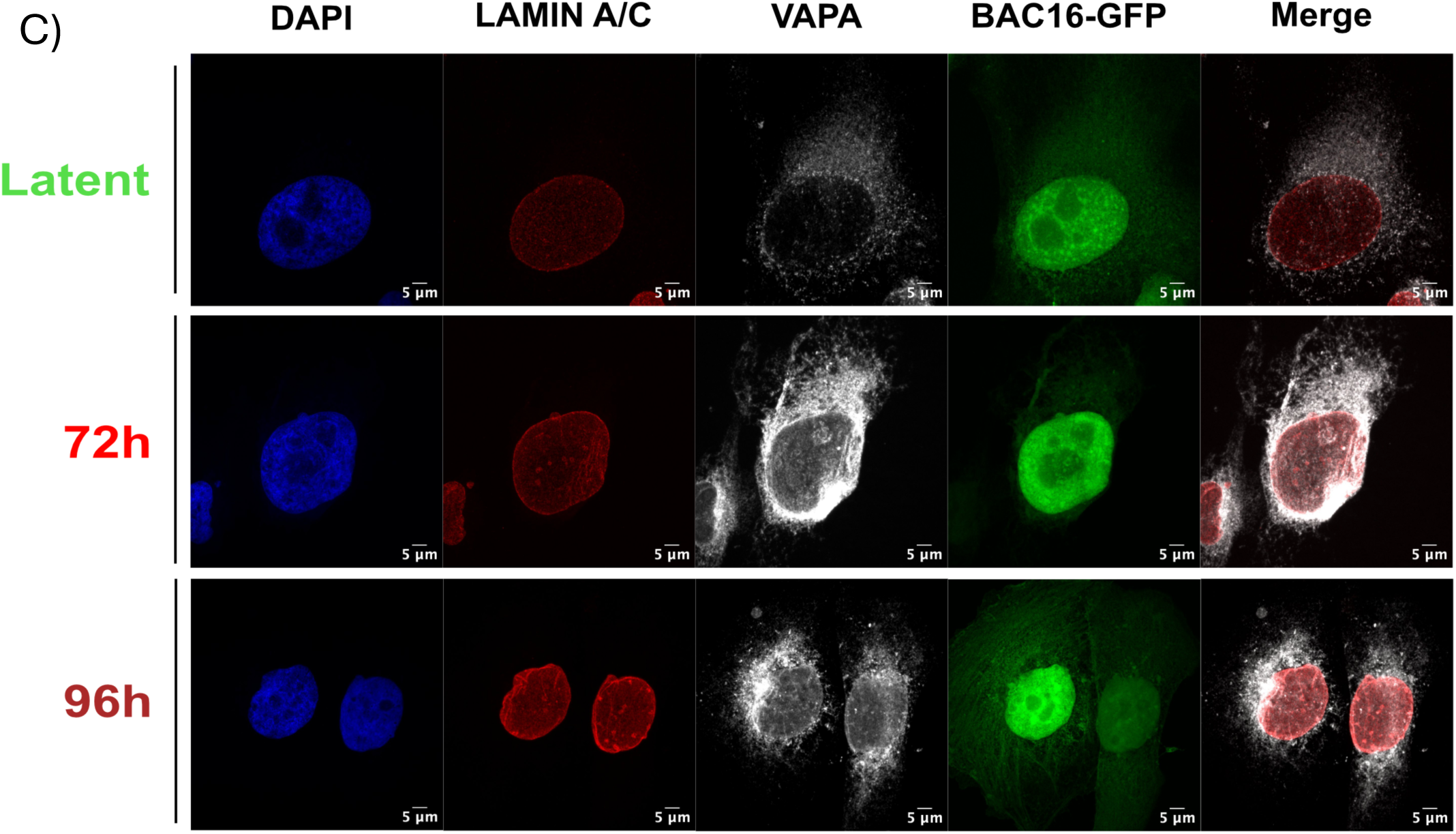
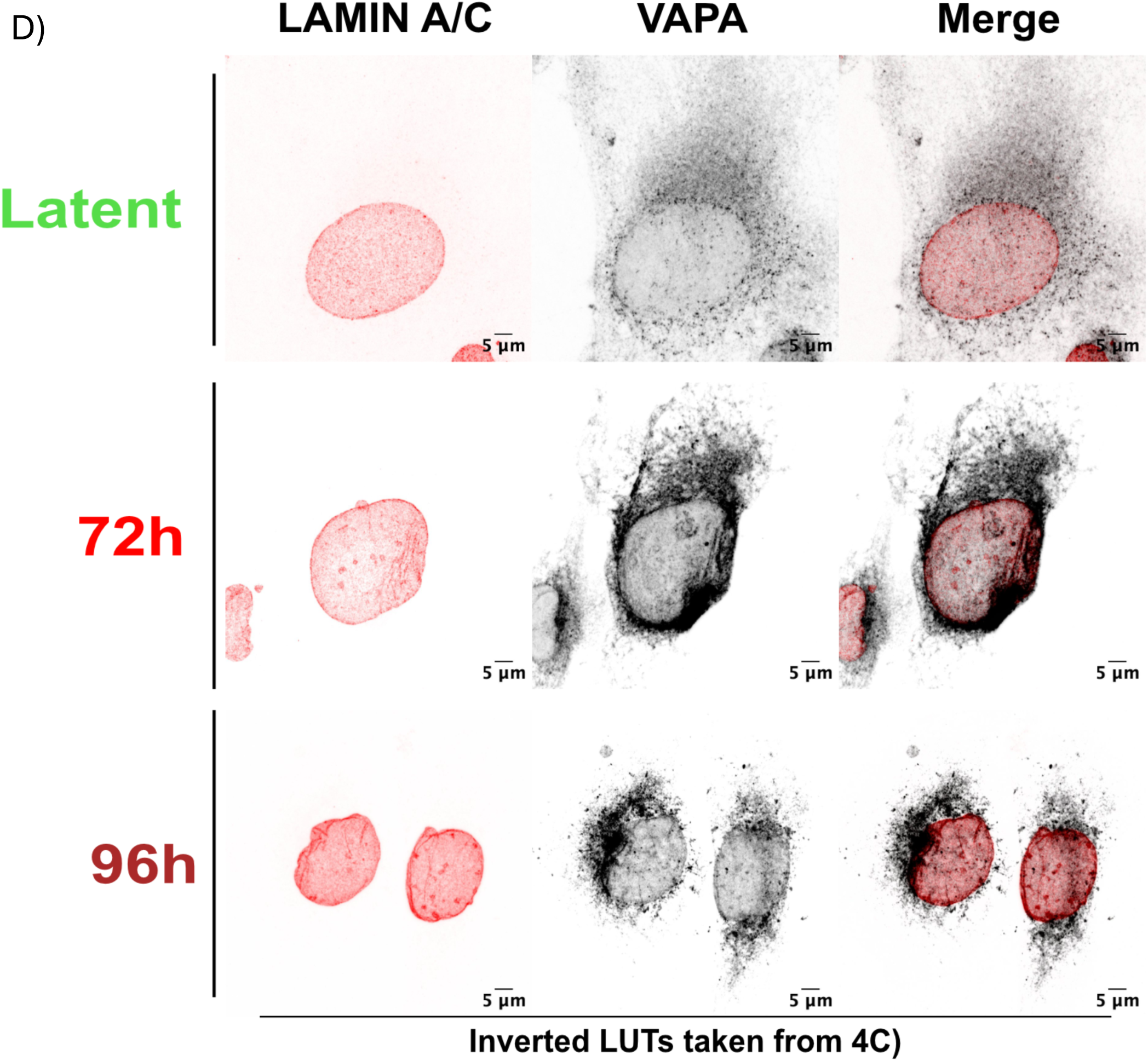
Type-I nucleoplasmic reticulum expansion in KSHV infected cells correlates with recruitment of CTP:phosphocholine cytidylyltransferase. (A) Doxycycline-inducible iSLK-BAC16 cells cultured on coverslips were reactivated from latency at 72 and 96h. Coverslips were fixed and immunostained for Lamin A/C and CCTα (A) or Lamin A/C and VAPA (C). Nuclei were counterstained with Hoechst and images were captured using a Zeiss LSM 880 confocal microscope (100× oil immersion). (B, D) Corresponding images from (A) and (C) with contrast enhancement and inverted lookup tables (LUTs) to better visualize intranuclear structures.

CCTα staining in latently KSHV infected cells was uniformly distributed in the nucleoplasm, with only occasional puncta, suggesting that CCTα primarily remains in its inactive state. By 72 and 96h post-lytic reactivation, CCTα puncta were prominent, indicating activation at membranes within the nucleoplasm. Fig. 5B (inverted LUT) shows CCTα puncta partially overlapping with lamin A/C-positive structures; this suggests that CCTα localizes to compartments partially covered in lamin A/C as well as those that are lamin A/C poor. VAPA staining in latent cells was primarily localized to the peripheral ER, with dim puncta scattered throughout the nucleus. At 72h, VAPA-positive veins and hollow loops appear near the nuclear periphery, characteristic of Type-II NR. By 96h, bright VAPA-positive puncta and tubules extend through the nucleus (Fig. 5C). Inverted LUT colorization (Fig. 5D) highlights co-localization of lamin A/C and VAPA, showing VAPA-positive tubules aligning with lamin A/C tubules. However, not all lamin A/C structures are VAPA-positive, indicating the presence of both Type-I and Type-II NR within these cells.

## Discussion

In herpesvirus egress, capsids bud into INM with the help of a two-component viral nuclear egress complex (NEC), briefly acquiring an envelope that is quickly shed via fusion with the ONM. Upon gaining access to the cytoplasm, these capsids bud into the *trans* Golgi network and travel to the cell surface in secretory vesicles that fuse with the plasma membrane to release progeny herpesviruses into the extracellular environment. The primary envelopment event at the INM has been observed at the nuclear periphery, but also at nuclear infolds (NIs) that provide a tantalizing route for ‘picking up’ newly minted capsids from viral replication compartments in the interior of the nucleus. However, mechanisms governing herpesvirus access to NIs remain poorly characterized. Here, we report that both Type-I and Type-II nucleoplasmic reticulum (NR) infolds increase during KSHV lytic replication, correlating with the recruitment of CCTα to these membranes, which drives membrane proliferation via phosphatidyl choline synthesis (Figure 6). There is emerging evidence that herpesvirus nuclear egress operates with a quality-control mechanism wherein intact DNA-containing C-capsids are selectively exported, leaving behind immature or defective capsids. Our study reveals another aspect of selectivity in KSHV nuclear egress; even though both Type-I and Type-II NR increase during KSHV lytic replication, we only observe budding into Type-I NR, along with accumulation de-enveloped C-capsids at 2^nd^ order infoldings. This suggests an attractive complementary mechanism of KSHV nuclear egress, whereby the Type-I NR invagination accesses new KSHV capsids at assembly sites, reducing the need for capsid trafficking to the peripheral INM, followed by capsid budding and fusion at the NR prior to release into the cytoplasm. Our static images of capsid accumulation at Type-I NR suggest that capsid budding and fusion at these sites is more rapid than the pace of NR retraction and capsid release. Future studies of these dynamic processes will benefit from the use of live cell imaging approaches with fluorescently labelled NR and capsids.

**Figure 6.**
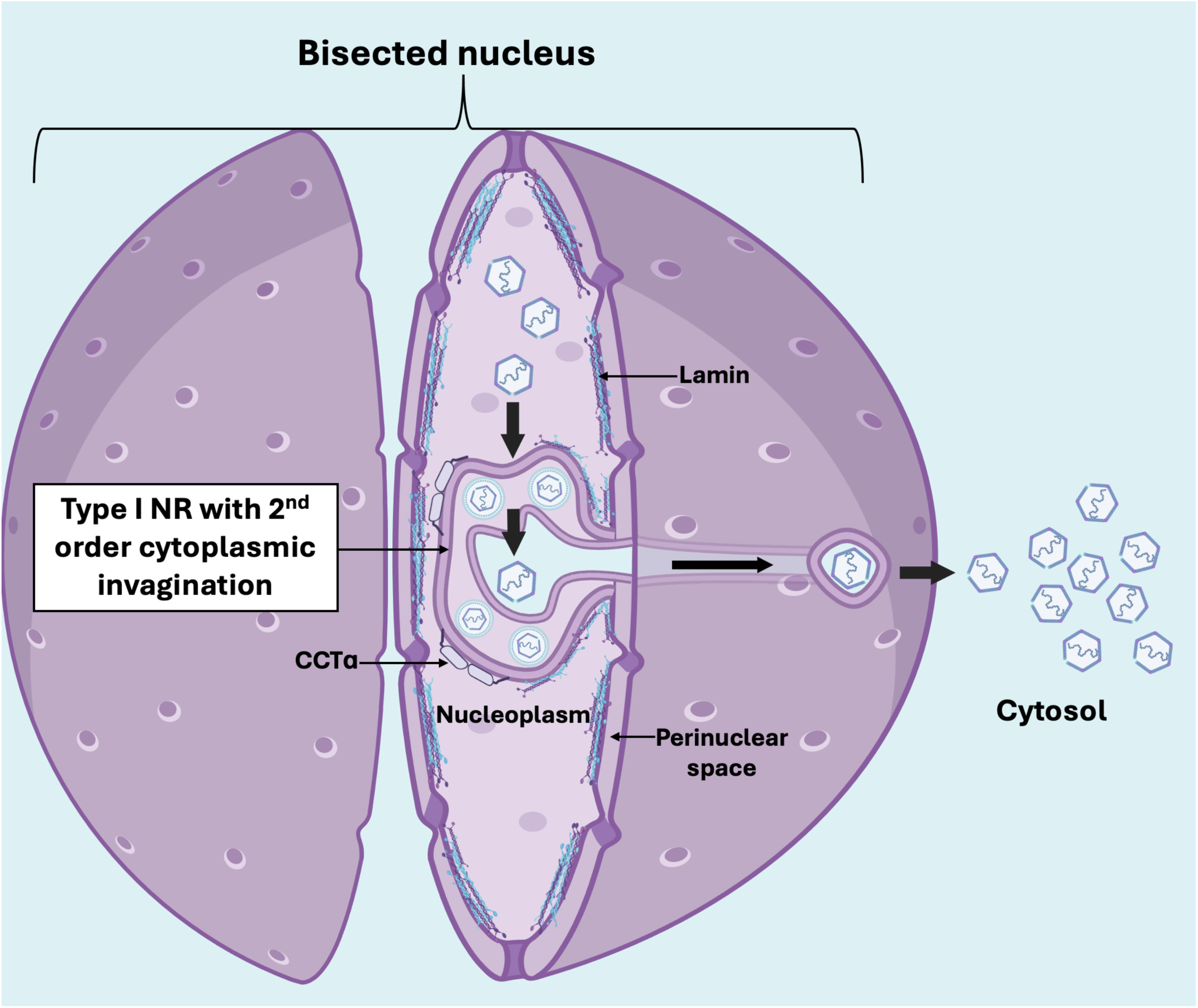
A model for KSHV primary envelopment at Type-I NR. The NR expands over the duration of lytic replication accompanied by a change in the distribution of lamin and CCTα. C-capsids acquire an envelope upon budding into the Type-I NR and exist as enveloped C-capsids within a Type-I NR-derived compartment that resembles a 1^st^ order nuclear infold (NI). Enveloped C-capsids lose their envelope upon budding out of the 1^st^ order NI into a compartment resembling a 2^nd^ order NI. These 2^nd^ order NIs are continuous with the cytoplasm enabling capsids to travel out of the 2^nd^ order structures and access the cytosol.

We observed KSHV budding at the Type-I NR and the peripheral INM in the same cell, suggesting that both mechanisms can operate concurrently. There were substantially more C-capsids associated with Type-I NR than peripheral INM in these images, suggesting that the NR could be a preferred route for nuclear egress that allows bulk transport of capsids rather than one-by-one at the peripheral INM. While these observations are largely consistent with previous studies of human alphaherpesviruses and betaherpesviruses, as well as murine gamma-herpesviruses, suggesting a conserved mechanism for accessing NR for primary envelopment, we do not yet understand the factors that influence which route of primary envelopment a capsid pursues. We also do not yet understand what factors might prevent capsid budding into Type-II NR, or why capsid de-envelopment occurs at 2^nd^ order infolds within Type-I NR, but not at greatly expanded convoluted membrane structures. These are important questions for future research.

In this paper we describe a novel structure we term the convoluted membrane NR (Figure 2B and 3A/B). We predict these membranes originate by mass expansion of the INM causing the membrane to fold into a structure resembling a membrane whorl. We observed bright CCTα positive puncta at some spherical compartments positively stained for lamin A/C (Figure 5A). We speculate that CCTα could preferentially localize or drive expansion of the INM and become concentrated in the convoluted membranes. Notably, CCTα can dimerize in *trans* (56), which may stabilize two opposing convoluted INMs. We suggest within the newly forming convoluted membrane, CCTα promotes the synthesis of PtdCho, promoting unchecked INM expansion and formation of convoluted structures.

Lamin forms a structural barrier around the inner nuclear membrane (INM), preventing herpesvirus capsids and proteins from accessing it. Data presented in Figure 5 demonstrates profound reorganization of lamin A/C during late lytic replication. The pattern of lamin redistribution during KSHV infection mirrors the lamin A/C staining during infection with HCMV (53), EBV (42), HSV-1 (55), and HSV-2 (54). Since herpesvirus capsids cannot bud through intact lamin, these viruses have evolved mechanisms to disrupt this barrier, facilitating primary envelopment at the INM. As noted in the Introduction, herpesviruses from all subfamilies encode viral proteins that reorganize the nuclear lamina during lytic replication. KSHV encodes a single serine/threonine protein kinase called ORF36, which is a conserved protein among other herpesviruses (57). ORF36 is expressed during early lytic replication and localizes to the nucleus. The KSHV tegument protein ORF45 engages ribosomal S6 kinase (RSK) and regulates the phosphorylation and activity of ORF36 (58). The same study also found that ORF45/RSK regulate the phosphorylation of lamin A. Thus, multiple KSHV proteins collaborate to remodel nuclear lamina to aid nuclear egress, although our understanding of these processes remains incomplete.

A strength of our findings is the extensive dataset of 2D TEM images that capture the morphology of Type-I NR during KSHV infection. Future studies employing 3D imaging, similar to the approach used for HCMV (19), could offer deeper insights into the number of C-capsids packed within these structures and the full extent of the nuclear protrusions. Supporting this, our IF data suggests a more extensive network than what is captured in a single 2D TEM section (Compare Fig. 2b with Fig. 5a/c). Overall, our findings provide substantial support for the proliferation of NR during lytic herpesvirus replication and the viral utilization of Type-I NR specifically as a route for nuclear egress. This should motivate future studies of the molecular mechanisms of NR expansion in infected cells and high-resolution studies of NR dynamics in living cells.

## Materials and methods

### Cell culture and reagents

Doxycycline inducible iSLK-BAC16 cells were cultured in Dulbecco’s modified Eagle’s medium (DMEM) (Gibco) supplemented with 10% fetal bovine serum, 2 mM L-glutamine, 1% penicillin-streptomycin and grown in 5% CO_2_ at 37°C. To maintain stability of episomal KSHV DNA in iSLK cells, parental cell cultures were maintained in 1 μg/ml puromycin (ThermoFisher) and 1 mg/ml hygromycin B (Invitrogen). iSLK-BAC16 cells were reactivated from latency with the addition of 1 mM sodium butyrate (Sigma) and 1μg/mL doxycycline (Sigma).

### Transmission electron microscopy

iSLK-BAC16 cells were seeded in 10 cm dishes (VWR) at a density of approximately 1.5 × 10^5^ cells per dish (5% CO_2_ at 37°C). The next day, cells were reactivated from latency by the addition of 1 mM sodium butyrate and 1 μg/mL doxycycline and harvested at 24, 48, 72 or 96 h. Cells were harvested by trypsin digestion and centrifuged at 250 x g for 5 minutes. Samples were fixed for a minimum of 2 hours in 2.5% glutaraldehyde diluted in 0.1 M sodium cacodylate buffer. Following fixation, samples were rinsed three times for at least 10 minutes each with 0.1M sodium cacodylate buffer. Secondary fixation was performed using 1% osmium tetroxide for 2 hours, followed by a brief rinse with distilled water. Samples were then placed in 0.25% uranyl acetate at 4°C overnight. Dehydration was carried out using a graded acetone series: 50% acetone for 10 minutes, 70% acetone for two 10-minute incubations, 95% acetone for two 10-minute incubations, and 100% acetone for two 10-minute incubations, followed by a final 10-minute incubation in dried 100% acetone. Samples were then infiltrated with Epon Araldite resin using a stepwise approach: 3:1 (dried 100% acetone to resin) for 3 hours, 1:3 (dried 100% acetone to resin) overnight, and finally 100% resin for two 3-hour incubations. Samples were embedded in 100% Epon Araldite resin and cured at 60°C for 48 hours. Ultrathin sections (∼100 nm thick) were obtained using a Reichert–Jung Ultracut E ultramicrotome equipped with a diamond knife and were placed on 300-mesh copper grids. Sections were stained with 2% aqueous uranyl acetate for 10 minutes, followed by two 5-minute rinses with distilled water. Lead citrate staining was performed for 4 minutes, followed by a quick rinse with distilled water, and the grids were air-dried. Prepared samples were imaged using a JEOL JEM 1230 transmission electron microscope operating at 80 kV. Images were acquired using a Hamamatsu ORCA-HR digital camera.

### Immunofluorescence and Image Processing

Doxycycline-inducible iSLK-BAC16 cells were seeded at 300,000 cells/mL on glass coverslips (Paul Marienfeld Gmb H & Co.; 0117580) in 12-well plates (VWR). The next day, cells were reactivated with 1 μg/mL doxycycline and 1mM sodium butyrate and harvested at 72h or 96 h. Coverslips were fixed in 4% paraformaldehyde for 15 min at room temperature, washed with PBS, and incubated in blocking/permeabilization (block/perm) buffer (Triton-X100 0.1% (v/v) (Sigma; 1002614889) and 1% human AB serum (Sigma) in PBS for 1 h at RT. Coverslips were incubated with mouse anti-Lamin A/C (1:200, Sigma) and/or rabbit anti-CCTα (1:500) and/or rabbit anti-VAPA (1:200) antibodies overnight at 4°C in block/perm buffer. The next day, coverslips were washed three times with PBS followed by a 1 h incubation at room temperature with secondary donkey anti-rabbit AlexaFluor555 (Invitrogen; A31572) and chicken anti-mouse AlexaFluor647 (Invitrogen; A21463) for CCTα immunofluorescence experiments, and goat anti-mouse AlexaFluor555 (Invitrogen; A21422) and goat anti-rabbit AlexaFluor647 (Invitrogen; A21244) for VAPA immunofluorescence. All secondary antibodies were diluted 1:1000 in block/perm buffer. After three additional PBS washes, coverslips were counterstained with Hoechst 33342 (1:5000, 5 min, RT) and mounted on microscope slides (Fisherbrand; 12-550-15) using ProLong Gold Antifade (ThermoFisher, P36930).

### Imaging & Data Processing

Z-stacks were acquired using a Zeiss LSM 880 fluorescence microscope (100× oil objective) and processed into maximum intensity projections using Zeiss Black software. Images were further analyzed in FIJI (ImageJ), with a global contrast increase of 0.34% for VAPA experiments. Inverted LUT images (Figure 4) were contrast-adjusted to highlight intranuclear structures. Notably, 96h samples exhibited intense CCTα staining, suggesting upregulation or aggregation. To compensate, the A568 laser power was reduced from 4 to ∼2.6 when imaging 96h samples.

### Data Transparency

All experiments were performed in three biological replicates, with multiple images of different cells included in the main text and supplementary data. To ensure transparency, additional TEM and IF images are available in our online data repository (LINK).

## Acknowledgements

We would like to thank Jae Jung and Soowon Kang for sending us their doxycycline-inducible iSLK-BAC16 cell line and providing their insights into the handling and culture of these cells. We also thank Mary Ann Trevors chief electron microscopy technologist within the CORES facility at Dalhousie University. Mary Ann provided invaluable support with sample preparation for TEM analysis alongside invaluable teaching and insight into interpreting TEM images.

This work has been supported by the Scotia Scholars Doctoral award.

## References

1. Tandon R, Mocarski ES, Conway JF. 2015. The A, B, Cs of herpesvirus capsids. Viruses 7:899–914.

2. Nealon K, Newcomb WW, Pray TR, Craik CS, Brown JC, Kedes DH. 2001. Lytic Replication of Kaposi’s Sarcoma-Associated Herpesvirus Results in the Formation of Multiple Capsid Species: Isolation and Molecular Characterization of A, B, and C Capsids from a Gammaherpesvirus. J Virol 75:2866–2878.

3. Wu L, Lo P, Yu X, Stoops JK, Forghani B, Zhou ZH. 2000. Three-Dimensional Structure of the Human Herpesvirus 8 Capsid. J Virol 74:9646–9654.

4. Costa RH, Cohen G, Eisenberg R, Long D, Wagner’ E, Frink J, Draper KG, Wagner EK. 1984. Direct demonstration that the abundant 6-kilobase herpes simplex virus type 1 mRNA mapping between 0.23 and 0.27 map units encodes the major capsid protein VP5. J Virol 49:287.

5. Lo P, Yu X, Atanasov I, Chandran B, Zhou ZH. 2003. Three-Dimensional Localization of pORF65 in Kaposi’s Sarcoma-Associated Herpesvirus Capsid. J Virol 77:4291.

6. Booy FP, Trus BL, Newcomb WW, Brown JC, Conway JF, Steven AC. 1994. Finding a needle in a haystack: detection of a small protein (the 12-kDa VP26) in a large complex (the 200-MDa capsid of herpes simplex virus). Proc Natl Acad Sci U S A 91:5652.

7. Rixon FJ, Davison MD, Davison AJ. 1990. Identification of the genes encoding two capsid proteins of herpes simplex virus type 1 by direct amino acid sequencing. J Gen Virol 71:1211–1214.

8. Okoye ME, Sexton GL, Huang E, McCaffery JM, Desai P. 2006. Functional Analysis of the Triplex Proteins (VP19C and VP23) of Herpes Simplex Virus Type 1. J Virol 80:929.

9. Newcomb WW, Juhas RM, Thomsen DR, Homa FL, Burch AD, Weller SK, Brown JC. 2001. The UL6 Gene Product Forms the Portal for Entry of DNA into the Herpes Simplex Virus Capsid. J Virol 75:10923.

10. Tsurumi S, Watanabe T, Iwaisako Y, Suzuki Y, Nakano T, Fujimuro M. 2021. Kaposi’s sarcoma-associated herpesvirus ORF17 plays a key role in capsid maturation. Virology 558:76–85.

11. Preston VG, Al-Kobaisi MF, McDougall IM, Rixon FJ. 1994. The herpes simplex virus gene UL26 proteinase in the presence of the UL26.5 gene product promotes the formation of scaffold-like structures. J Gen Virol 75:2355–2366.

12. Stevens A, Kashyap S, Crofut E, Lucia Alverez-Cabrera A, Jih J, Liu Y-T, Zhou H. Structure of a new capsid form and comparison with A-, B- and C-capsids clarify herpesvirus assembly 10.1101/2025.03.19.644230.

13. Panté N, Kann M. 2002. Nuclear pore complex is able to transport macromolecules with diameters of ∼39 nm. Mol Biol Cell 13:425–434.

14. Desai PJ, Pryce EN, Henson BW, Luitweiler EM, Cothran J. 2012. Reconstitution of the Kaposi’s Sarcoma-Associated Herpesvirus Nuclear Egress Complex and Formation of Nuclear Membrane Vesicles by Coexpression of ORF67 and ORF69 Gene Products. J Virol 86:594–598.

15. Reynolds AE, Ryckman BJ, Baines JD, Zhou Y, Liang L, Roller RJ. 2001. UL31 and UL34 Proteins of Herpes Simplex Virus Type 1 Form a Complex That Accumulates at the Nuclear Rim and Is Required for Envelopment of Nucleocapsids. J Virol 75:8803.

16. Milbradt J, Kraut A, Hutterer C, Sonntag E, Schmeiser C, Ferro M, Wagner S, Lenac T, Claus C, Pinkert S, Hamilton ST, Rawlinson WD, Sticht H, Couté Y, Marschall M. 2014. Proteomic Analysis of the Multimeric Nuclear Egress Complex of Human Cytomegalovirus. Mol Cell Proteomics 13:2132–2146.

17. Roffman E, Albert JP, Goff JP, Frenkel1 N. 1990. Putative site for the acquisition of human herpesvirus 6 virion tegument. J Virol 64:6308.

18. Buser C, Walther P, Mertens T, Michel D. 2007. Cytomegalovirus Primary Envelopment Occurs at Large Infoldings of the Inner Nuclear Membrane. J Virol 81:3042–3048.

19. Villinger C, Neusser G, Kranz C, Walther P, Mertens T. 2015. 3D Analysis of HCMV Induced-Nuclear Membrane Structures by FIB/SEM Tomography: Insight into an Unprecedented Membrane Morphology. Viruses 7:5686–5704.

20. Peng L, Ryazantsev S, Sun R, Zhou ZH. 2010. Three-Dimensional Visualization of Gammaherpesvirus Life Cycle in Host Cells by Electron Tomography. Structure 18:47–58.

21. McPhee M, Dellaire G, Ridgway ND. 2024. Mechanisms for assembly of the nucleoplasmic reticulum. Cell Mol Life Sci C 81:415.

22. Ruebner BH, Miyai K, Slusser RJ, Wedemeyer P, Medearis DN. 1964. Mouse Cytomegalovirus Infection: An Electron Microscopic Study of Hepatic Parenchymal Cells. Am J Pathol 44:799.

23. Papadimitriou JM, Shellam GR, Robertson TA. 1984. An ultrastructural investigation of cytomegalovirus replication in murine hepatocytes. J Gen Virol 65:1979–1990.

24. Nassiri MR, Gilloteaux J, Taichman RS, Drach JC. 1998. Ultrastructural aspects of cytomegalovirus-infected fibroblastic stromal cells of human bone marrow. Tissue Cell 30:398–406.

25. Dal Monte P, Pignatelli S, Zini N, Maraldi NM, Perret E, Prevost MC, Landini MP. 2002. Analysis of intracellular and intraviral localization of the human cytomegalovirus UL53 protein. J Gen Virol 83:1005–1012.

26. Malhas A, Goulbourne C, Vaux DJ. 2011. The nucleoplasmic reticulum: form and function. Trends Cell Biol 21:362–373.

27. Sutter E, de Oliveira AP, Tobler K, Schraner EM, Sonda S, Kaech A, Lucas MS, Ackermann M, Wild P. 2012. Herpes simplex virus 1 induces de novo phospholipid synthesis. Virology 429:124–135.

28. Fuchs W, Klupp BG, Granzow H, Osterrieder N, Mettenleiter TC. 2002. The Interacting UL31 and UL34 Gene Products of Pseudorabies Virus Are Involved in Egress from the Host-Cell Nucleus and Represent Components of Primary Enveloped but Not Mature Virions. J Virol 76:364.

29. Lorenz M, Vollmer B, Unsay JD, Klupp BG, García-Sáez AJ, Mettenleiter TC, Antonin W. 2015. A Single Herpesvirus Protein Can Mediate Vesicle Formation in the Nuclear Envelope. J Biol Chem 290:6962.

30. Klupp BG, Granzow H, Fuchs W, Keil GM, Finke S, Mettenleiter TC. 2007. Vesicle formation from the nuclear membrane is induced by coexpression of two conserved herpesvirus proteins. Proc Natl Acad Sci U S A 104:7241–7246.

31. Hagen C, Guttmann P, Klupp B, Werner S, Rehbein S, Mettenleiter TC, Schneider G, Grünewald K. 2012. Correlative VIS-fluorescence and soft X-ray cryo-microscopy/tomography of adherent cells. J Struct Biol 177:193.

32. Hagen C, Dent KC, Zeev-Ben-Mordehai T, Grange M, Bosse JB, Whittle C, Klupp BG, Siebert CA, Vasishtan D, Bäuerlein FJB, Cheleski J, Werner S, Guttmann P, Rehbein S, Henzler K, Demmerle J, Adler B, Koszinowski U, Schermelleh L, Schneider G, Enquist LW, Plitzko JM, Mettenleiter TC, Grünewald K. 2015. Structural Basis of Vesicle Formation at the Inner Nuclear Membrane. Cell 163:1692–1701.

33. Lv Y, Shen S, Xiang L, Jia X, Hou Y, Wang D, Deng H. 2019. Functional Identification and Characterization of the Nuclear Egress Complex of a Gammaherpesvirus. J Virol 93:e01422–19.

34. Reynolds AE, Ryckman BJ, Baines JD, Zhou Y, Liang L, Roller RJ. 2001. U L 31 and U L 34 Proteins of Herpes Simplex Virus Type 1 Form a Complex That Accumulates at the Nuclear Rim and Is Required for Envelopment of Nucleocapsids. J Virol 75:8803–8817.

35. Scott ES, O’Hare P. 2001. Fate of the Inner Nuclear Membrane Protein Lamin B Receptor and Nuclear Lamins in Herpes Simplex Virus Type 1 Infection. J Virol 75:8818–8830.

36. Mou F, Forest T, Baines JD. 2007. US3 of Herpes Simplex Virus Type 1 Encodes a Promiscuous Protein Kinase That Phosphorylates and Alters Localization of Lamin A/C in Infected Cells. J Virol 81:6459.

37. Leach NR, Roller RJ. 2010. Significance of host cell kinases in herpes simplex virus type 1 egress and lamin-associated protein disassembly from the nuclear lamina. Virology 406:127–137.

38. Park R, Baines JD. 2006. Herpes Simplex Virus Type 1 Infection Induces Activation and Recruitment of Protein Kinase C to the Nuclear Membrane and Increased Phosphorylation of Lamin B. J Virol 80:494–504.

39. Wu S, Pan S, Zhang L, Baines J, Roller R, Ames J, Yang M, Wang J, Chen D, Liu Y, Zhang C, Cao Y, He B. 2016. Herpes Simplex Virus 1 Induces Phosphorylation and Reorganization of Lamin A/C through the γ 1 34.5 Protein That Facilitates Nuclear Egress. J Virol 90:10414–10422.

40. Bjerke SL, Roller RJ. 2006. Roles for herpes simplex virus type 1 UL34 and US3 proteins in disrupting the nuclear lamina during herpes simplex virus type 1 egress. Virology 347:261–276.

41. Cano-Monreal GL, Wylie KM, Cao F, Tavis JE, Morrison LA. 2009. Herpes simplex virus 2 UL13 protein kinase disrupts nuclear lamins. Virology 392:137.

42. Lee C-P, Huang Y-H, Lin S-F, Chang Y, Chang Y-H, Takada K, Chen M-R. 2008. Epstein-Barr Virus BGLF4 Kinase Induces Disassembly of the Nuclear Lamina To Facilitate Virion Production. J Virol 82:11913–11926.

43. Sharma M, Kamil JP, Coughlin M, Reim NI, Coen DM. 2014. Human Cytomegalovirus UL50 and UL53 Recruit Viral Protein Kinase UL97, Not Protein Kinase C, for Disruption of Nuclear Lamina and Nuclear Egress in Infected Cells. J Virol 88:249–262.

44. Hamirally S, Kamil JP, Ndassa-Colday YM, Lin AJ, Jahng WJ, Baek MC, Noton S, Silva LA, Simpson-Holley M, Knipe DM, Golan DE, Marto JA, Coen DM. 2009. Viral Mimicry of Cdc2/Cyclin-Dependent Kinase 1 Mediates Disruption of Nuclear Lamina during Human Cytomegalovirus Nuclear Egress. PLoS Pathog 5:e1000275.

45. Brulois KF, Chang H, Lee AS-Y, Ensser A, Wong L-Y, Toth Z, Lee SH, Lee H-R, Myoung J, Ganem D, Oh T-K, Kim JF, Gao S-J, Jung JU. 2012. Construction and Manipulation of a New Kaposi’s Sarcoma-Associated Herpesvirus Bacterial Artificial Chromosome Clone. J Virol 86:9708–9720.

46. Drozdz MM, Jiang H, Pytowski L, Grovenor C, Vaux DJ. 2017. Formation of a nucleoplasmic reticulum requires de novo assembly of nascent phospholipids and shows preferential incorporation of nascent lamins. Sci Reports 2017 71 7:1–14.

47. Pytowski L, Drozdz MM, Jiang H, Hernandez Z, Kumar K, Knott E, Vaux DJ. 2019. Nucleoplasmic Reticulum Formation in Human Endometrial Cells is Steroid Hormone Responsive and Recruits Nascent Components. Int J Mol Sci 2019, Vol 20, Page 5839 20:5839.

48. Iwaisako Y, Watanabe T, Futo M, Okabe R, Sekine Y, Suzuki Y, Nakano T, Fujimuro M, Jung JU. 2022. The Contribution of Kaposi’s Sarcoma-Associated Herpesvirus ORF7 and Its Zinc-Finger Motif to Viral Genome Cleavage and Capsid Formation. J Virol 96.

49. Santos MF, Rappa G, Karbanová J, Kurth T, Corbeil D, Lorico A. 2018. VAMP-associated protein-A and oxysterol-binding protein–related protein 3 promote the entry of late endosomes into the nucleoplasmic reticulum. J Biol Chem 293:13834–13848.

50. Kent C. 1997. CTP:phosphocholine cytidylyltransferase. Biochim Biophys Acta - Lipids Lipid Metab 1348:79–90.

51. Gehrig K, Cornell RB, Ridgway ND. 2008. Expansion of the Nucleoplasmic Reticulum Requires the Coordinated Activity of Lamins and CTP:Phosphocholine Cytidylyltransferase. Mol Biol Cell 19:237–247.

52. Lagace TA, Ridgway ND. 2005. The Rate-limiting Enzyme in Phosphatidylcholine Synthesis Regulates Proliferation of the Nucleoplasmic Reticulum. Mol Biol Cell 16:1120–1130.

53. Camozzi D, Pignatelli S, Valvo C, Lattanzi G, Capanni C, Dal Monte P, Landini MP. 2008. Remodelling of the nuclear lamina during human cytomegalovirus infection: Role of the viral proteins pUL50 and pUL53. J Gen Virol 89:731–740.

54. Cano-Monreal GL, Wylie KM, Cao F, Tavis JE, Morrison LA. 2009. Herpes simplex virus 2 UL13 protein kinase disrupts nuclear lamins. Virology 392:137–147.

55. Simpson-Holley M, Colgrove RC, Nalepa G, Harper JW, Knipe DM. 2005. Identification and Functional Evaluation of Cellular and Viral Factors Involved in the Alteration of Nuclear Architecture during Herpes Simplex Virus 1 Infection. J Virol 79:12840–12851.

56. Taneva SG, Patty PJ, Frisken BJ, Cornell RB. 2005. CTP:phosphocholine cytidylyltransferase binds anionic phospholipid vesicles in a cross-bridging mode. Biochemistry 44:9382–9393.

57. Park J, Lee D, Seo T, Chung J, Choe J. 2000. Kaposi’s sarcoma-associated herpesvirus (human herpesvirus-8) open reading frame 36 protein is a serine protein kinase. J Gen Virol 81:1067–1071.

58. Avey D, Tepper S, Li W, Turpin Z, Zhu F. 2015. Phosphoproteomic Analysis of KSHV-Infected Cells Reveals Roles of ORF45-Activated RSK during Lytic Replication. PLOS Pathog 11:e1004993.

59. Pignatelli S, Dal Monte P, Landini MP, Severi B, Nassiri R, Gilloteaux J, Papadimitriou JM, Shellam GR. 2007. Cytomegalovirus Primary Envelopment at Large Nuclear Membrane Infoldings: What’s New? J Virol 81:7320.

